# Integration of scHi-C and scRNA-seq data defines distinct 3D-regulated and biological-context dependent cell subpopulations

**DOI:** 10.1101/2023.09.29.560193

**Authors:** Yufan Zhou, Tian Li, Lavanya Choppavarapu, Victor X. Jin

**Author notes:** Correspondence (V.X.J).

## Abstract

An integration of 3D chromatin structure and gene expression at single-cell resolution has yet been demonstrated. Here, we develop a computational method, a multiomic data integration (MUDI) algorithm, which integrates scHi-C and scRNA-seq data to precisely define the 3D-regulated and biological-context dependent cell subpopulations or topologically integrated subpopulations (TISPs). We demonstrate its algorithmic utility on the publicly available and newly generated scHi-C and scRNA-seq data. We then test and apply MUDI in a breast cancer cell model system to demonstrate its biological-context dependent utility. We found the newly defined topologically conserved associating domain (CAD) is the characteristic single-cell 3D chromatin structure and better characterizes chromatin domains in single-cell resolution. We further identify 20 TISPs uniquely characterizing 3D-regulated breast cancer cellular states. We reveal two of TISPs are remarkably resemble to high cycling breast cancer persister cells and chromatin modifying enzymes might be functional regulators to drive the alteration of the 3D chromatin structures. Our comprehensive integration of scHi-C and scRNA-seq data in cancer cells at single-cell resolution provides mechanistic insights into 3D-regulated heterogeneity of developing drug-tolerant cancer cells.

## Introduction

Three-dimension (3D) chromatin architecture within a nucleus can be constructed from chromosome conformation capture (3C) related techniques including 3C^1^, 4C^2^, 5C^3^, ChIA-PET^4^, Hi-C^5^, TCC^6^ and in situ Hi-C^7^. These profiling methods have revealed major 3D genomic features, including genomic compartments^5,8^, topologically associating domains (TADs)^9^ and chromatin loops^7^. Many computational methods have been simultaneously developed to determine these features, including normalizing interacting contact maps^8,10^, computing A/B compartments^5,11^, calling TADs^12,13^, detecting significant interactions^7,14,15^, enhancing the low sequencing depth data^16,17^, and visualizing the contact matrices^18–21^. Further, in order to delineate the heterogeneity of population cells, single-cell Hi-C (scHi-C) protocols have been newly developed to identify 3D chromatin architecture at single-cell resolution^22–25^. For instance, the dynamic chromosomal organization of cell cycle^26^, the organization of zygote chromatin^27,28^, the nuclear changes of stem cell differentiation^29^, and single-allele chromatin interactions^30,31^ have been fully examined by scHi-C technique. Meanwhile, new sets of computational methods have been developed for processing scHi-C data to reconstruct single-cell 3D chromatin^32–34^, to impute the chromosome contact matrices^35–37^, to identify TAD-like domains^38^, to classify single cells^39^, to identify chromatin loops^40^, and to provide toolbox of scHi-C^41^. However, none of these methods were designed to algorithmically integrate scHi-C and single-cell (sc)RNA-seq data. Therefore, it is imperative to develop a method for comprehensively integrating single-cell chromatin domains and single-cell gene expression to precisely define 3D-regulated cell subpopulations.

Drug-tolerant cancer cells (DTCCs) are a subpopulation of cancer cells that resist the anti-cancer drug treatment and likely cause the patient relapse after therapeutics. DTCCs usually consists of three different groups according to the period of drug treatment^42^. The first group is cancer persister cells survived in the short-term drug shock. The second group is extended persister cells revived and proliferated in the mid-term drug stress. The third group is stable drug-resistant cancer cells survived with clonal selection in the long-term drug treatment. Studies have shown that genetic^43^ or non-genetic mechanisms^44,45^ were involved in regulating the development of DTCCs. In our recent study, we found that the dynamic changes of 3D chromatin structures might be a non-genetic mechanism driving breast cancer endocrine resistance^46^. However, the patterning and characteristics of 3D chromatin structures in DTCCs at single-cell resolution have not been elucidated.

Here, we develop a computational method, a multiomic data integration (MUDI) algorithm, which integrates scHi-C and scRNA-seq data to precisely define the 3D-regulated and biological-context dependent cell subpopulations or topologically integrated subpopulations (TISPs). We demonstrate its algorithmic utility on the publicly available and newly generated scHi-C and scRNA-seq data. We then apply MUDI in a breast cancer cell model system, including three stages of breast cancer cells, tamoxifen-sensitive breast cancer cells (MCF7), MCF7 cells after being temporally treated with tamoxifen for one month (MCF7M1), and MCF7 derived tamoxifen-resistant cells (MCF7TR) after being temporally treated with tamoxifen for six months. We identify and characterize distinct 3D-regulated cancer cell subpopulations, and further determine 3D-regulated heterogeneity of developing drug-tolerant cancer cells.

## Results

### Developing a computational method to integrate scHi-C and scRNA-seq data

To comprehensively integrate scHi-C and scRNA-seq data, we developed a novel computational method, a multiomic data integration (MUDI) algorithm, to precisely define 3D-regulated cell subpopulations or TISPs (Fig. 1a). We first identified distinct scHi-C clusters from scHi-C data, and scRNA-seq clusters from scRNA-seq data, respectively. We then integrated these two types of clusters by the MUDI algorithm (see Methods: Integration of scHi-C and scRNA-seq data) to precisely define the distinct TISPs (Fig. 1a). Briefly, we first defined topologically conserved associating domains (CADs) representing the conserved 3D chromatin structure of any individual scHi-C cluster. We then integrated CADs with differentially expressed genes (DEGs) of each of scRNA-seq clusters to derive TISPs by implementing an empirical quantitative formula to calculate an integration score of the interaction frequency and the gene expression values. We tested our MUDI on two cell types: pluripotent stem cells WTC11 from 4D Nucleome Project of Bing Ren Lab and breast cancer cells MCF7 generated from this study. From scHi-C data, nine scHi-C clusters (CC1-CC9) were identified with variable relative contact probability (Fig. 1b-c, Extended Data Fig. 1a-c), where CC1/3/5/7 and CC2/4/6/8/9 are majorly composed of WTC11 cells and MCF7 cells, respectively. From scRNA-seq data, ten scRNA-seq clusters (DD1-DD10) were classified with variable fold changes of differentially expressed genes (DEGs) (Fig. 1d-e). DD1/2/4/5/7/8/9 and DD3/6/10 are majorly composed of WTC11 cells and MCF7 cells, respectively. Our MUDI was initially able to identify four TISPs (WMG1-WMG4) with the distinct subpopulation features based on the number (M) of data types (here M = 2) and the number of (N) of cell types (here N = 2), such that WMG1 is the subpopulation with integration of CC1/3/5/7 and DD1/2/4/5/7/8/9, WMG2 is the subpopulation with the integration of CC1/3/5/7 and DD3/6/10, WMG3 is the subpopulation with integration of CC2/4/6/8/9 and DD1/2/4/5/7/8/9, WMG4 is the subpopulation with integration of CC2/4/6/8/9 and DD3/6/10 (Extended Data Fig. 1d-e). More importantly, the MUDI is further designed to be tailored to a biological-context dependent integration, such that the number of TISPs can be optimized according to a particular biologically meaningful factor on individual studies. Since Yamanaka Factors, MYC, POU5F1, SOX2, KLF4, were used to characterize the stem cell differentiation, we were able to obtain 12 distinct TISPs (Fig. 1f, Extended Data Fig. 2a), where one of subpopulations YFG1 was enriched with REACTOME developmental biology signaling pathway (Extended Data Fig. 2b-c), suggesting this subpopulation has high stemness and strong chromatin activities.

**Fig. 1.**
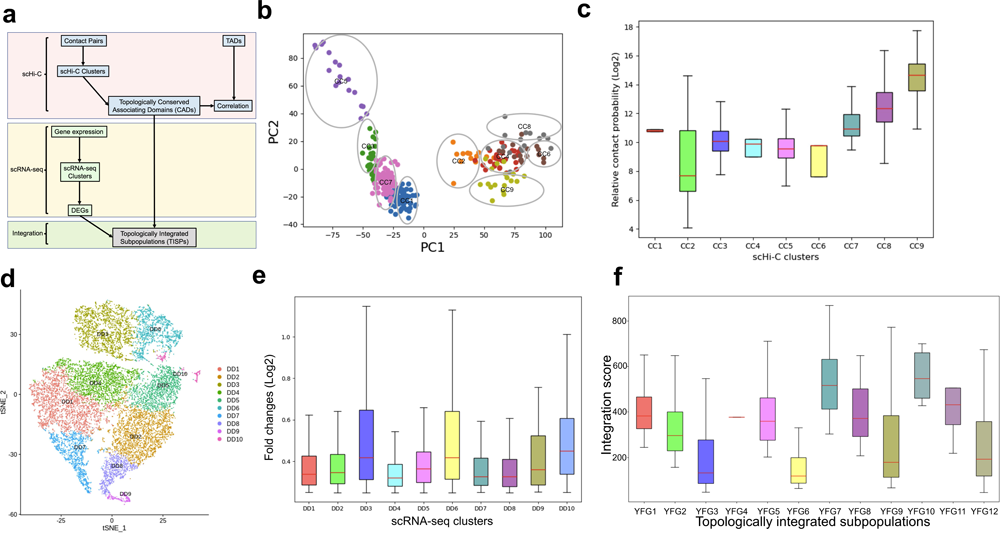
Development of a computational method for integrating scHi-C and scRNA-seq data. **a**, Flowchart of Multiomic Data Integration (MUDI) algorithm. DEGs: Differentially expressed genes. TADs: Topologically associating domains. **b**, Nine scHi-C clusters (CC1-CC9) identified from scHi-C data of WTC11/MCF7. **c**, Relative contact probability of scHi-C clusters. **d**, Ten scRNA-seq clusters (DD1-DD10) identified from scRNA-seq data of WTC11/MCF7. **e**, Fold changes of DEGs of scRNA-seq clusters. **g**, Integration scores of 12 topologically integrated subpopulations (TISPs), YFG1-12.

To further demonstrate the sensitivity and robustness of the MUDI, we have first performed a sub-sampling analysis on WTC11 cells and MCF7 cells (Extended Data Fig. 3a). We found that compared to the whole set of 277 cells, it showed no significant difference of the overlapped CADs in each cluster for the subset of 75% (208) cells and the subset of 50% (138) cells, respectively, but significant difference for the subset of less than 25% (69) cells. Therefore, our MUDI algorithm is sensitive to at least half of cells. We then tested the MUDI on sn-m3c-seq data generated from human brain tissues^87^. After the integration of human cortex sn-m3c-seq data (Extended Data Fig. 3b) and human cortex scRNA-seq data (Extended Data Fig. 3c)^88^, we identified TISP specifying excitatory neurons with a set of 55 cell markers of excitatory neurons (Extended Data Fig. 3d). These genes formed interaction networks and the excitatory neuron cell marker SATB2 was one of network hubs (Extended Data Fig. 3e). Furthermore, our MUDI was successfully applied in three datasets with significantly different sequencing depths, including 1) sn-m3C-seq data of human prefrontal cortex tissue with an average of 1.2M contact pairs per cell, 2) scHi-C data of WTC11 cells with an average of 10.5M contact pairs per cell, and 3) our newly generated scHi-C data of three breast cancer cells with an average of 36.4M contact pairs per cell (see next four sections). Our MUDI has been able to identify computationally significant and biologically meaningful TISPs, suggesting that our algorithm was much less dependent on the sequencing depth. In summary, we have developed a novel and powerful method, MUDI, to precisely define 3D-regulated and biological-context dependent cell subpopulations.

### Generating high quality scHi-C and scRNA-seq data in a breast cancer cell model system

In order to further test and demonstrate the biological-context dependent utility of MUDI, we have generated high quality scHi-C and scRNA-seq data in a breast cancer cell model system, MCF7, MCF7M1 and MCF7TR cells (Fig. 2a), a model system routinely used in the lab^46^. A total of 293 cells (89 MCF7 cells, 91 MCF7M1 cells, 113 MCF7TR cells) were used for scHi-C profiling (Extended Data Fig. 4a) and 22,425 cells (6,172 MCF7 cells, 10,156 MCF7M1 cells, 6,097 MCF7TR cells) were used for scRNA-seq profiling (Extended Data Fig. 4b). Single cell chromatin contacts with very high quality were obtained (Extended Data Fig. 4c) upon preprocessing scHi-C data (Extended Data Fig. 4d-e, Extended Table 1), The combined scHi-C data showed a significant correlation with population Hi-C data, *i.e.*, correlation coefficient r = 0.43 for combined single cells MCF7 to population MCF7, r = 0.61 for combined single cells MCF7M1 to population MCF7M1, and r = 0.58 for combined single cells MCF7TR to population MCF7TR, respectively. The correlations were weak among combined single cells, *i.e.*, correlation coefficient r = 0.05 for combined single cells MCF7 to combined single cells MCF7M1, r = 0.28 for combined single cells MCF7M1 to combined single cells MCF7TR, r = 0.07 for combined scHi-C MCF7 to combined scHi-C MCF7TR, respectively (Fig. 2b). Genomic distance dependent contact probability showed markedly characteristic shapes of combined single cells (Fig. 2c, upper left) and individual single cells (Fig. 2c, upper right, lower left, and lower right panels). We also observed that the single cells had highly variable TADs but with more superimposing of cells, the enriched TADs have more similar features of population TADs (Fig. 2d-f). These results demonstrated a high quality of scHi-C data had been successfully produced in cancer cells. Since single-cell omics-seq data are generally sparse, an optimal resolution is needed for the downstream analysis. Our scHi-C data have a low slope of ratio of read pairs to square of bin numbers until the resolution reaches to 1Mb (Extended Data Fig. 4f), thus the 1Mb resolution was used for clustering of scHi-C data.

**Fig. 2.**
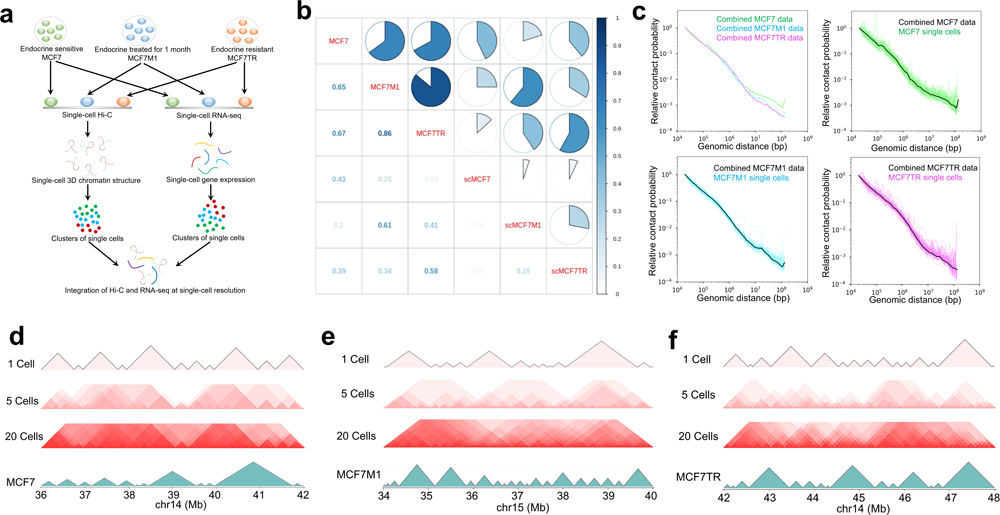
Generation of high quality scHi-C and scRNA-seq data in a breast cancer cell model system. **a**, Workflow for the identification of 3D chromatin structures of breast cancer cell lines at single-cell resolution. **b**, Pearson correlation coefficients of combined scHi-C data with population data. scMCF7: combined MCF7 scHi-C data. scMCF7M1: combined MCF7M1 scHi-C data. scMCF7TR: combined MCF7TR scHi-C data. **c**, Genomic distance dependent contact probability. The thick lines are combined single cells and the thin lines are individual single cells. **d-f**, Superimposing single-cell TADs with 5 or 20 cells compared to population Hi-C TADs. (**d**) for MCF7, (**e**) for MCF7M1, and (**f**) for MCF7TR, respectively. All TADs were generated at the resolution of 100Kb contact map.

To exclude the effect of structure variations (SVs), we performed single-cell DNA-seq on three breast cancer cell lines each with a biological replicate: 33 MCF7 cells, 33 MCF7M1 cells and 39 MCFTR cells with a total of 105 cells. We found that 1) there was no clear difference on copy number variations (CNVs) among single cells (Extended Data Fig. 4g), 2) scHi-C contacts in the genomic regions where 10% cells had CNVs had a very low ratio (almost zero) and 3) there was not any significant difference between MCF7 cells and MCF7TR cells (Extended Data Fig. 4h). These results illustrated that single-cell level SVs didn’t significantly influence the chromatin contacts.

### Defining the characteristic single-cell 3D chromatin structure

Before performing scHi-C clustering, we first examined our scHi-C data quality by comparing it with publicly available human scHi-C data. The breast cancer cells from our study were clearly separated from other types of human cells, leukemia cells K562^27^ and two pluripotent stem cell types, WTC11C6 and WTC11C28 (4D Nucleome Project, Bing Ren Lab) (Fig. 3a, Extended Data Fig. 5a-b). Furthermore, three stages of breast cancer cells, MCF7, MCF7M1 and MCF7TR were also distinctly located in different spaces defined by first three eigenvectors (Fig.3b-c, Extended Data Fig. 5c). This analysis further validated the high quality of our scHi-C data. We then applied scHiCluster^36^ to identify an optimal nine scHi-C clusters, C1 to C9 (Fig. 3d) since the peak of the Sihouette coefficient is at 9 (Extended Data Fig. 5d). We removed the cells with the contacts lower than 6 in 1Mb bins to minimize the false positive rate (Extended Data Fig. 6a-d) and thus obtained a good quality of 231 cells (87 MCF7 cells, 54 MCF7M1 cells and 90 MCFTR cells). Of nine clusters, a majority of cells in C2 and C7 were MCF7, a majority of cells in C1, C3, C4, C8, C9 were MCF7TR, and the cells in C5 and C6 were miscellaneous of three stages of cells (Fig. 3e). Interestingly, C1 and C5 had the smallest size of TADs and the most numbers of TADs (Fig. 3f, Extended Data Fig. 7a), while MCF7M1 cells had smaller sizes of TADs than MCF7 and MCF7TR cells did (Extended Data Fig. 7b-c).

**Fig. 3.**
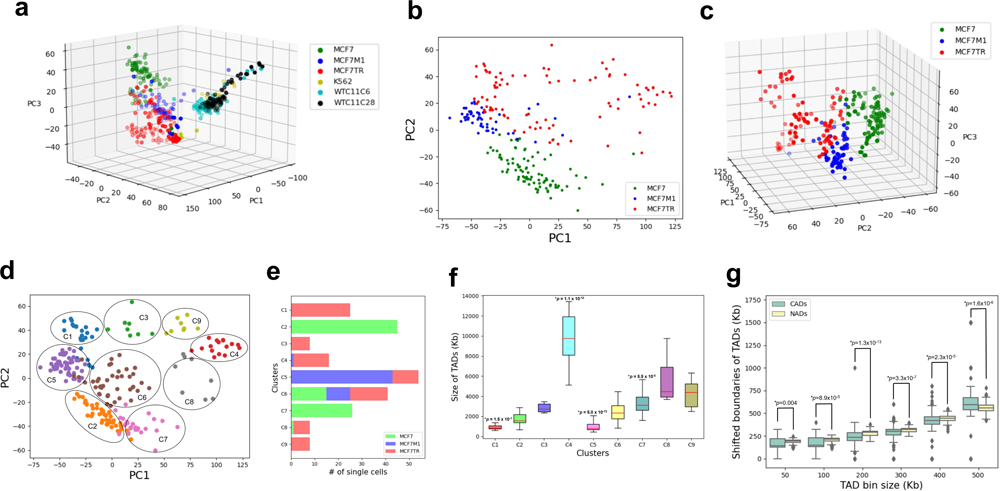
Definition of the characteristic single-cell chromatin structure. **a**, Comparing our scHi-C data with public human scHi-C data. PC1, PC2 and PC3 are first three eigenvectors. **b**, 2D view of scHi-C data of breast cancer cells. **c**, 3D view of scHi-C data of breast cancer cells. **d**, Nine clusters (C1-C9) identified from scHi-C data of breast cancer cells. Each cluster is labelled with oval and assorted colors. **e**, Number and the composition of single cells in individual scHi-C clusters. **f**, The size of TADs of clusters. *: Wilcoxon rank-sum test. **g**, The shifted boundaries of TADs of CADs and NADs when TAD bin size is 50K, 100K, 200K, 300K, 400K or 500K. *: Wilcoxon rank-sum test.

To better characterize chromatin domains in single-cell resolution, We proposed a novel framework for analyzing 3D chromatin domain behavior among single cells, and defined a CAD which is the common 1Mb genomic region shared by all individual cells within any particular scHi-C cluster that has very high chromatin contact probabilities. Indeed, CADs showed lower shifted boundaries of TADs and greater standard deviations than non-conserved associating domains (NADs) (Fig. 3g, Extended Data Fig. 8a). CADs had different characteristics from NADs in each of nine clusters. For example, CADs in C1 showed the highest shifted boundaries in compared to NADs at 100Kb TAD size (Extended Data Fig. 8b-d, 9a-f), and there were the most CADs either in all cells or per cell for C1, C3, C5, and C9 (Extended Data Fig. 7d-e). Our results thus elucidated that the newly defined CAD is the characteristic single-cell 3D chromatin structure useful for functional analysis of scHi-C clusters.

### Precisely identifying distinct 3D-regulated cancer cell subpopulations

To precisely identify the 3D-regulated cancer cell subpopulations, we further conducted scRNA-seq data (Extended Data Fig. 10a-b) with the replicates showing a highly identical pattern in MCF7, MCF7M1 and MCF7TR cells (Extended Data Fig. 10c). We then identified 13 scRNA-seq clusters, D1-D13 (Fig. 4a), in which a majority of cells in D2, D6, and D11 are MCF7, a majority of cells in D1, D4, D5, D8, D9 and D10 are MCF7M1, a majority of cells in D3, D7, D12, D13 are MCF7TR (Fig. 4b). We also identified a gene signature of differentially expressed genes (DEGs) for each of 13 clusters (Fig. 4c, Extended Table 2). Interestingly, we found that the cell cycle signaling was among the top enriched pathways from the top 2000 variably expressed genes (Extended Data Fig. 10d, 11a) and the standardized variance of cycling genes is much higher than that of housekeeping genes (Fig. 4d, Extended Table 3). More specifically, there were much more cycling genes within DEGs in D3, D5, D7, D8, D10 as well as within CADs in C1, C3, C5, C9 than other scHi-C or scRNA-seq clusters (Fig. 4e). Remarkably, cycling signaling has been used to characterize cancer persister cells, a rare subpopulation of DTCCs with a reversible property^45^. We thus grouped scHi-C clusters into five categories based on the breast cancer cell stage and the number (high: >9; low: =<9) of cycling genes within CADs: 1) C1, C5 --miscellaneous cells with high cycling genes; 2) C6 -- miscellaneous cells with low cycling genes; 3) C3, C9 – resistant cells with high cycling genes; 4) C4, C8 -- resistant cells with low cycling genes; 5) C2, C7 -- sensitive cells with low cycling genes. Miscellaneous cells either with high cycling genes (C1, C5) or with low cycling genes (C6) showed higher contact probabilities than sensitive cells (C2, C7) (Extended Data Fig. 11b-c). On the contrary, resistant cells regardless of with high (C3, C9) or low (C4, C8) cycling genes had lower contact probabilities than sensitive cells (C2, C7) (Extended Data Fig. 10d-e). Although both Categories 1) and 3) have high cycling genes, miscellaneous cells (C1, C5) have more contact probabilities than resistant cells (C3, C9) (Extended Data Fig. 11f). We then computed an integration score within MUDI program to integrate five scHi-C categories with four scRNA-seq categories, and thus precisely defined 20 TISPs, G1-20, each representing a 3D-regulated breast cancer cellular state by an integration score (Fig. 4f).

**Fig. 4.**
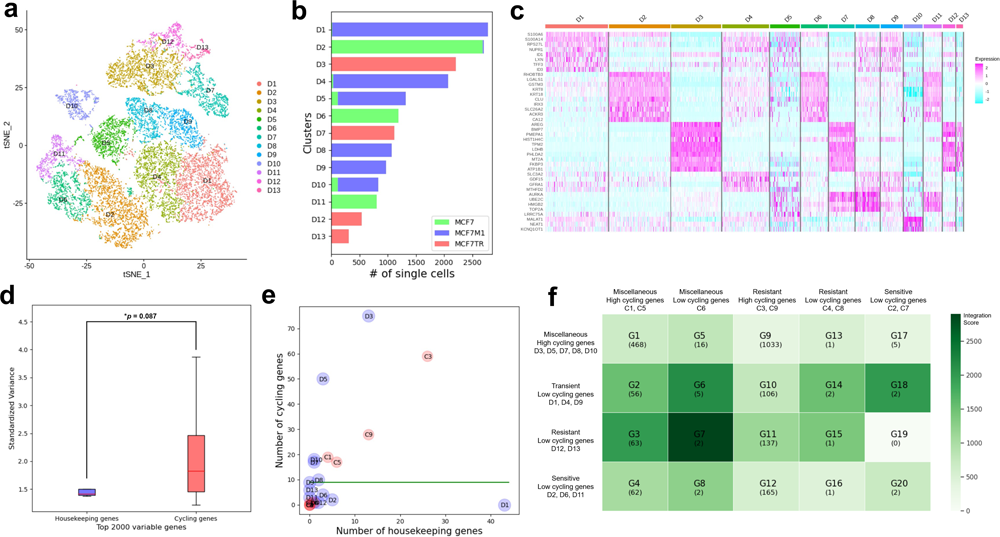
Precise identification of 3D-regulated and biological-context dependent cancer cell subpopulations. **a**, Thirteen scRNA-seq clusters (D1-D13) identified from scRNA-seq data of breast cancer cells. **b**, Number and the composition of single cells in individual scRNA-seq clusters. **c**, Gene expression heatmap of DEGs of scRNA-seq clusters. **d**, The standardized variance of cycling genes and housekeeping genes in top 2000 variable genes. *: Wilcoxon rank-sum test. **e**, The distribution of CADs in scHi-C clusters and DEGs in scRNA-seq clusters according to the number of cycling genes and the number of housekeeping genes in each cluster. Green line is the cutoff for high cycling genes and low cycling genes. **f**, Twenty topologically integrating subpopulations (TISPs) (G1-G20) dependent on the number of cycling genes and cell compositions of the scHi-C clusters and scRNA-seq clusters.

### Characterizing specific topologically integrated subpopulations

We further examined a few of the TISPs related to cycling genes. Despite both G1 and G9 had high cycling genes in both CADs of scHi-C clusters and DEGs of scRNA-seq clusters, G1 had a higher integration score than G9 (Fig. 5a, Extended Data Fig. 12a). In addition, some of G1 and G9 genes were marked with super-enhancers (Extended Data Fig. 12b-c). Interestingly, G1 genes were enriched with a REACTOME chromatin modifying enzyme signaling pathway and these enriched enzymes had higher integration scores in G1 than those in G9 (Fig. 5b-c). Of 15 enriched genes, ATXN7, ENY2, PRMT6, KDM5B, KMT5A, MBIP, SMARCB1, TADA3 occurred in G1 and G9, BRWD1, CCND1, ELP2, HMG20B, JADE1, KMT2E, MORF4L1 in G9 (Extended Data Fig. 12d). Higher expression of chromatin modifying enzymes in breast cancer patient cohorts showed a lower recurrence-free survival (Fig. 5d, Extended Data Fig. 12e-k). Of these genes, CCND1, ENY2 and KMT5A had epithelial cell-specific *cis*-regulatory elements at their distal regions in luminal breast cancer patient tissue^89^. Together, these results suggest G1 and G9 might resemble to cycling breast cancer persister cells and their 3D chromatin structures might be regulated by chromatin modifying enzymes.

**Fig. 5.**
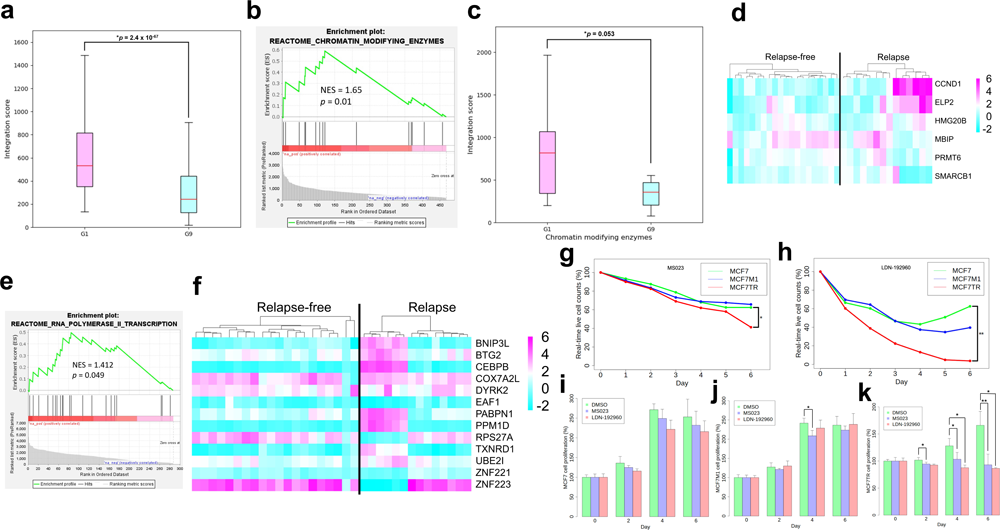
Characteristics of TISPs in breast cancer cells. **a**, The integration score of G1 and G9. *: Wilcoxon rank-sum test. **b**, Enrichment of REACTOME chromatin modifying enzymes signaling pathway of G1 genes. NES: Normalized Enrichment Score. **c**, Comparison of the integration score between G1 and G9. *: Wilcoxon rank-sum test. **d**, The expression of chromatin modifying enzymes in relapse-free and relapse breast cancer patient cohort GSE2990. **e**, Enrichment of REACTOME RNA polymerase II transcription signaling pathway of the combination of G2, G3, G10 and G11. NES: Normalized Enrichment Score. **f**, The expression of transcription regulators in relapse-free and relapse breast cancer patient cohort GSE2990. **g**, Real-time live cell growth curve of PRMT6 inhibitor MS023. Cells treated with DMSO as reference. *: *p* < 0.05, paired Student’s t-test. **h**, Real-time live cell growth curve of DYRK2 inhibitor LDN-192960. Cells treated with DMSO as reference. **: *p* < 0.01, paired Student’s t-test. **i**, Cell proliferation assay of MS023 and LDN-192960 in MCF7 cells. **j**, Cell proliferation assay of MS023 and LDN-192960 in MCF7M1 cells. *: *p* < 0.05, paired Student’s t-test. **k**, Cell proliferation assay of MS023 and LDN-192960 in MCF7TR cells. *: *p* < 0.05, **: *p* < 0.01, paired Student’s t-test.

On the other hand, cell subpopulations, G2, G3, G10 and G11, had high cycling genes in CADs of scHi-C clusters but low cycling genes in DEGs of scRNA-seq clusters. REACTOME RNA polymerase II transcription signaling pathway was the top enriched pathway from these four subpopulations (Fig. 5e). Of 21 enriched genes, CEBPB and YEATS4 existed in G2, THOC7 and TXNRD1 in G2 and G10, and COX7A2L, RPS27A, UBE2I, ZNF221 and ZNF223 in G10, while RPRD1A existed in G3, NELFA, PPM1D and SRAF1 in G3 and G10, and BNIP3L, BTG2, CNOT6, DYRK2, EAF1, MED1, PABPN1 and TIGAR in G10 (Extended Data Fig. 13a). Higher expression of transcription regulators in breast cancer patient cohorts was correlated with a lower recurrence-free survival (Fig. 5f, Extended Data Figs. 13b-h, 14a-e). Among them, CEBPB, COX7A2L, NELFA, SRSF1, TXNRD1, UBE2I had epithelial cell specific *cis*-regulatory elements at their distal regions in luminal breast cancer patient tissue^89^. Collectively, these results suggest that these four cell subpopulations might resemble to non-cycling breast cancer persister cells and their 3D chromatin structures might be regulated by transcription regulators.

To further substantiate our findings, we performed an experimental validation for the drug treatment on the two selected genes identified by our MUDI, PRMT6 and DYRK2. The section of these two genes was purely due to the commercially available inhibitors to them. We treated MS023, an inhibitor to PRMT6, a key regulator in G1 and G9 subpopulations, and LDN-192960, an inhibitor to DYRK2, a key transcriptional regulator in G10. We found both inhibitors showed stronger growth inhibition in MCF7TR cells than that in MCF7 cells (Fig. 5g,h), as well as impeded MCF7TR cells from cell proliferation but not MCF7 (Fig. 5i-k), demonstrating the capability of the inhibitors of these regulators in restoring the drug-sensitivity.

Taken together, we propose a mechanistic model with two distinct 3D-regulated cellular states for the transition of drug-sensitive to tolerant cancer cells: 1) a drug-sensitive cancer cell subpopulation with silenced chromatin modifying enzymes initially shows very lower chromatin interactions (Extended Data Fig. 15a); upon an interim drug treatment, this subpopulation activates the enzymes to trigger higher chromatin interacting activities for the cycling genes, resulting in reversible cancer persister cells (Extended Data Fig. 15b); under a long-term drug treatment, they further reshape the altered 3D chromatin structures render a cycling drug-tolerant cancer cells (Extended Data Fig. 15c); and 2) another drug-sensitive cancer cell subpopulation with silenced transcription regulators initially shows lower chromatin interactions (Extended Data Fig. 15d); upon an interim drug treatment, this subpopulation activates transcription regulators to trigger higher chromatin interacting activities for the non-cycling genes, resulting in reversible cancer persister cells (Extended Data Fig. 15e); under a long-term drug treatment, they further reshape the altered 3D chromatin structures render a non-cycling drug-tolerant cancer cells (Extended Data Fig. 15f).

## Discussion

In this study, we developed a novel computational method, MUDI, to comprehensively integrate scHi-C and scRNA-seq data and to precisely define distinct 3D-regulated and biological-context dependent cell subpopulations or TISPs. In the MUDI, we first defined CADs representing the conserved 3D chromatin structure of any individual scHi-C cluster. We then integrated CADs with DEGs of each of scRNA-seq clusters to derive TISPs by implementing an empirical quantitative formula to calculate an integration score of the interaction frequency and the gene expression values. A high integration score of a TISP indicates it is strongly associated with a set of higher expressed genes with higher chromatin interacting activities. More importantly, the identified TISPs are readily used to interpret biological-context dependent 3D-regulated cell subpopulations according to a particular biologically meaningful factor on individual studies. Furthermore, these 3D-regulated and biological-context dependent cell subpopulations can be used to elucidate a specific biological mechanism.

Remarkably, upon the application of MUDI in three stages of breast cancer cells, we illustrated cycling breast cancer cell subpopulations (miscellaneous or resistant) have distinctive altered 3D chromatin structures regulated by different regulators. It is reasonable to speculate these cell subpopulations resemble to breast cancer persister cells. Future studies will be focused on functionally examination of breast cancer persister cells. We may apply a Watermelon, a high-complexity expressed barcode lentiviral library^45^ to simultaneously trace each breast cancer Tam-sensitive cell’s clonal origin and proliferative state with a short period series of Tam-treatment (0-14 days), then conduct 3D-FISH, 3C/RT-qPCR and Tam-treatment to confirm if cycling persister cells is indeed 3D-regulated and can be re-sensitized.

Interestingly, we found that cell cycle genes highly enriched within CADs were a key factor to stratify the Tam-sensitive cells from 1-month Tam-treated and Tam-resistant cells. Indeed, many studies have demonstrated cell cycle pathway played important roles in breast cancer tamoxifen resistance^47–51^. For instance, cyclin D1 was essential for the progression of tamoxifen resistance^47^ and inner nuclear membrane protein LEM4 activated cell cycle proteins to render tamoxifen resistance^50^, Importantly, our data further linked cell cycle signaling with 3D chromatin organization. This finding is pretty novel but not very surprising given that our other recent studies have demonstrated 3D chromatin architecture was associated with endocrine resistance^46,52–54^.

Furthermore, we identified two key groups of genes, 15 chromatin modifying enzymes and 21 transcriptional regulators, which were not only essential in 3D-regulated breast cancer cellular states, but also predicted a lower recurrence-free survival. Many of these genes have been extensively demonstrated their functional or mechanistic roles in different cancers^55–69^. For example, Protein arginine methyltransferase PRMT6 was shown to advance the progression in gastric cancer^57^, endometrial cancer^58^ and lung cancer^59^. Transcription factor CEBPB stimulated the metabolic reprogramming to increase the occurrence of cancer^64^. Phosphorylation of transcription mediator MED1 increased the drug resistance in prostate cancer^67^.

Overall, we demonstrated 3D-regulated cancer cell subpopulations were distinctly associated with different functional regulators. Our work might provide mechanistic insights into 3D-regulated heterogeneity of developing drug-tolerant cancer cells, giving a rationale in designing novel therapeutics of treating drug-tolerant cancer.

## Methods

### MUDI algorithm

After identifying scHi-C clusters by scHiCluster^36^, and scRNA-seq clusters by Seurat^70^, the CADs of each scHi-C cluster were integrated with DEGs of each scRNA-seq cluster to acquire integration scores. We defined the integration score calculated by individual genes present both in CADs and DEGs as the following:

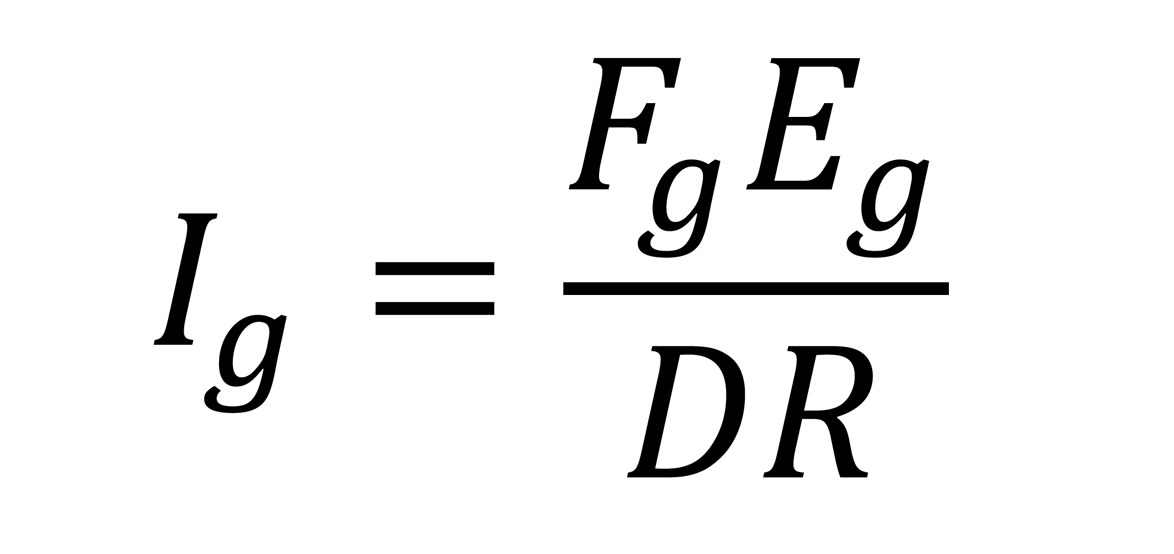

where I_g_ is the integration score of a gene. F_g_ is the relative contact probability (log_2_) of scHi-C data. E_g_ is expression fold changes (log_2_) of DEGs of scRNA-seq data. D is the ratio of DEGs of scRNA-seq clusters to total DEGs. R is the ratio of scRNA-seq cluster cells to total cells. “g” represents genes present in both scHi-C clusters and scRNA-seq clusters. The statistical *p* value of the difference of integration score was computed by Wilcoxon rank-sum test. We further classified scHi-C clusters into appropriate X scHi-C categories and scRNA-seq clusters into appropriate Y scRNA-seq categories by the biological-contexts, cell types or stages. Finally, product of X and Y is the total number of subpopulations. Each subpopulation has genes with integration score representing the expression level and chromatin interaction probability. The source code of MUDI is available at https://github.com/yufanzhouonline/MUDI.

### Data processing for scHi-C data

The raw reads of scHi-C were first aligned to human HG19 genome, then filtered by HiC-Pro version 2.11.1^71^ to get the valid pairs. The correlation of combined single cells to population cells was performed at the resolution of 1 Mb with R package HiCRep version 1.11.0^72^. The relative contact probabilities of individual cells were computed by cooltools version 0.4.0^73^ with the compensation of combined single cells. The TADs were called by Insulation Score^12^ at 100 Kb resolution if not specifically mentioned. The clustering of single cells was executed by Python package scHiCluster version 0.1.0^36^. Commonly Associating Domains (CADs) were defined as the common domains in a particular cluster at the resolution of 1 Mb, and non-commonly associating domains (NADs) were those non-common domains in that cluster. The difference of CADs, NADs and TADs was calculated with Wilcoxon rank-sum test. Super-enhancers were called with ChIP-seq data of H3K27ac in tamoxifen resistant MCF7 cells^46^ by Rank Ordering of Super-Enhancers (ROSE)^74^.

### Data processing for scRNA-seq data

The raw reads of scRNA-seq were first aligned to human HG19 genome and then feature-barcode matrices were generated with software Cell Ranger developed by 10X Genomics. The gene expression levels were further identified by Seurat version 4.0.3^70^ with the filtering parameters of min.cells at 3 and min.features at 200 on the module of CreateSeuratObject, and percent.mt < 30 on the module of subset. The resolution for finding clusters was set to 0.75 on the module of FindClusters. The differentially expressed genes (DEGs) of clusters were defined by the module of FindAllMarkers with the parameters of min.pct at 0.25 and logfc.threshold at 0.25. The difference of standardized variance between housekeeping genes and cycling genes in top 2000 variable genes were computed with Wilcoxon rank-sum test.

### Cell lines and reagents

Human breast cancer parental MCF7 cells and tamoxifen resistant MCF7TR cells were derived from previous study^46,75–77^. Temporal tamoxifen resistant MCF7M1 cells were generated from parental MCF7 cells treated with 100nM tamoxifen metabolite 4-hydroxytamoxifen (4-OHT) (Sigma, Catalog # H7904-5MG) for 1 month (30 days). MCF7, MCF7M1 and MCF7TR cells were cultured in phenol-free RPMI1640 medium (Thermo Fisher Scientific, Catalog # 11835055) supplemented with 10% charcoal stripped fetal bovine serum (FBS) (Sigma, Catalog # F6765-500ML) and 1% Penicillin-Streptomycin (Thermo Fisher Scientific, Catalog # 15140122), while no 4-OHT for MCF7 and MCF7M1 but supplemented with 100nM 4-OHT for MCF7TR.

### *In situ* Hi-C (population cells) profiling

In situ Hi-C experiments were performed as previously described with minor modifications^12^. Two to five million cells were crosslinked with 1% formaldehyde and then lysed with 0.2 Igepal CA630 to get the cell nuclei. The pelleted nuclei were solubilized with 0.5% sodium dodecyl (SDS) and then digested with restriction enzyme HindIII or DpnII. The restriction fragment overhangs were filled with biotin-14-dATP. The crosslinked proximity DNA was ligated with T4 DNA ligase. The crosslinked proteins were degraded by proteinase K. The DNA was pelleted down with ethanol and with sonication. A size of 300-500bp DNA was selected with AMPure XP beads and then the biotinylated DNA was pulled down with Dynabeads MyOne Streptavidin T1 beads. The ends of sheared DNA were repaired with DNA polymerase I. After the ligation of the adapter, the Hi-C libraries were amplified and purified. The libraries were sequenced on Illumina HiSeq 3000 Sequencer. Each sample was conducted in biological replicates. The sequencing reads were mapped to human HG19 genome with further normalization and filtering by HiC-Pro^71^.

### scHi-C profiling

Single-cell Hi-C experiment was performed majorly referring to Flyamer et al^27^ with minor revision. Two to four million MCF7 parental cells were fixed for 10 minutes by resuspending the cell pellet in 5 ml full culture medium supplemented with 1% formaldehyde. The reaction was quenched by addition of 2 M glycine to a final concentration of 125 mM and incubation for 5 min on ice. After washed with phosphate-buffered saline (PBS), cells were resuspended in lysis buffer (50mM Tris-HCl pH 8.0, 150 mM NaCl, 0.5% NP-40, 1% Triton X-100, 1X protease inhibitor cocktail and incubated on ice for at least 45 min. The lysed cell pellet was resuspended in 100 µl of 0.3% SDS in 1X NEBuffer 3 and incubated at 37°C for 1 hour. Then the resuspension was diluted with 330 µl of 1X NEBuffer 3 and 53 µl of 20% Triton X-100 and incubated at 37°C for 1 hour to quench SDS. The chromatin pellet was further digested with 600U restriction enzyme DpnII (New England BioLabs, Catalog # R0543M) overnight at 37°C with rotation. On the second day digestion was inactivated by incubation at 65°C for 20 min. The digested cell nuclei were ligated with 50U T4 DNA ligase for 4 hours and then washed with sterile PBS. The sample was stained with two drops of Hoechst 33342 (Thermo Fisher Scientific, Catalog # R37165) for 30 min at 37°C. Single cells were picked up by FACS sorter and loaded into 96 well PCR plate which each well filled with 5 µl sample buffer from the GenomiPhi V2 DNA amplification kit (previously GE Healthcare currently Cytiva, Catalog # 25660032), covered by 5 µl mineral oil after the sorting, then incubated at 65°C overnight. The genomic DNA were amplified according to Kumar et al^78^. The amplified genomic DNA of amounts more than 1 µg were prepared for sequencing with NEBNext Ultra II DNA Library Prep Kit for Illumina (New England BioLabs, Catalog # E7645L).

### scRNA-seq profiling

Cells were digested with 0.5% Trypsin-EDTA (Thermo Fisher Scientific, Catalog # 15400054) at the optimal time to avoid cell death and cell aggregation. After centrifugation, the cell pellet was resuspended in PBS (Thermo Fisher Scientific, Catalog # 14190250) at the concentration of 700-1200 cells per µl. If the viability of cells was higher than 90%, cells were then filtered with 40 µm sterile cell strainer (Fisher Scientific, Catalog # 22363547) to get individual cells. The samples of single cells were loaded on 10X Genomics Chromium system to run single-cell RNA-seq protocol according to the technical manual.

### scDNA-seq profiling

MCF7, MCF7M1 and MCF7TR cells were collected and sent to BioSkryb Genomics for isolation of single cell and scDNA-seq libraries preparation with the approach of Primary Template-directed Amplification (PTA)^85^. ResolveDNA Whole Genome Amplification Kit (Catalog # 100136, BioSkryb Genomics) was used for amplification of genomic DNA. ResolveDNA Library Preparation Kit (Catalog # 100080, BioSkryb Genomics) was used for the library construction. Libraries of scDNA-seq were sequenced on Illumina NovaSeq 6000 system. Sequencing raw reads were mapped to human HG19 genome and copy number variation was identified by SCCNV version 1.0.2^86^.

### Enrichment of signaling pathway

For scRNA-seq data, genes were pre-ranked by standardized variance then enriched by Gene Set Enrichment Analysis (GSEA) version 4.1.0^79^. Kyoto Encyclopedia of Genes and Genomes (KEGG) were used as gene sets database. For integrated scRNA-seq and scHi-C data, genes were pre-ranked by integration score then enriched by GSEA. REACTOME Pathway Database were used as gene sets database.

### Recurrence-free survival analysis

Two cohorts of breast cancer patients were used for survival analysis. Cohort GSE2990 was from Sotiriou et al^80^ and cohort GSE6532 was from Loi et al^81^. The patients were filtered by having tamoxifen treatment but no radio therapy or no other chemotherapy. The survival analysis was performed by R package Survival version 3.2-11. The patients were stratified by gene expression levels at the top quartile (25%) as high expression vs. the rest (75%) as low expression. The log-rank test was used for calculation of *p* value.

### Incucyte real-time live cell imaging

For a real-time live cell imaging of MCF7, MCF7M1 and MCF7TR, cells were seeded in 96-well plates at a density of 1 x 10^3^ cells per well. The cell media was replaced after 24 hrs and cells were treated with MS023 (10µm) and LDN (5µm) and the proliferation is monitored by the analysis of occupied area (% confluence) of cell images over time. As cells proliferate, the confluence increases. Confluence was an exceptional replacement for proliferation, until cells were densely packed or when large changes in morphology occurred. The graphs from the phase of cell confluence area were recorded from day 0 to day 6 according to the IncuCyte S3 Live-Cell Analysis System (Sartorius) manufacturer’s instructions. Incucyte S3 software version 2020B was used for the analysis.

### Cell proliferation assay

Cell viability was measured by CCK-8 (CCK-8, Dojindo, USA) assay following the manufacturer’s instructions. In brief, MCF7, MCF7M1 and MCF7TR cells were harvested and plated at a density of 1 x 10^3^ cells per well in 96-well plates (Corning Inc) and cultured in an incubator 5% CO_2_ incubator at 37°C. After 24 hrs, the culture media was replaced, and the cells are treated with MS023 (10µm) and LDN (5µm). At the end of each time point, 10 μL of CCK-8 solution was added to each 96-well plate and the mixture was incubated for 1 hr in the incubator at 37°C. The OD value of each well was measured by BioTek™ ELx800™ Absorbance Microplate Reader at 450nm. The assay was repeated three times.

### Simulation of 3D chromatin structure

Compartments of single cells were called by CscoreTool version 1.1^11^ at 50Kb resolution with the compensation of combined single cells. The compartments were then annotated as A1 (Cscore >= 0 and <= 0.2), A2 (Cscore > 0.2), B1 (Cscore < 0 and > -0.2) and B2 (Cscore <= -0.2) followed by simulation with chromatin dynamics software Open-MiChroM version 1.0.0^82^. The simulated structures were visualized by UCSF Chimera version 1.15^83^.

## Data availability

Raw and processed scHi-C data for MCF7, MCF7M1 and MCF7TR cells are deposited in GEO under accession number GSE194308. Raw and processed scRNA-seq data for MCF7, MCF7M1 and MCF7TR cells are deposited in GEO under accession number GSE195610, and raw and processed *in situ* Hi-C data for MCF7, MCF7M1 and MCF7TR cells are deposited in GEO under accession number GSE195810. Raw and processed scDNA-seq data for MCF7, MCF7M1 and MCF7TR cells are deposited in GEO under accession number GSE239435. WTC11C6 and WTC11C28 scHi-C datasets are publicly available datasets from 4D Nucleome Project Data Portal under accession number 4DNESJQ4RXY5 and 4DNESF829JOW. WTC11 scRNA-seq datasets are publicly available datasets from the ArrayExpress database under accession number E-MTAB-6268^84^.

## Author contributions

VXJ conceived the project. YZ conducted the experiments and performed the data analysis. TL and LC assisted in conducting the experiments. VXJ and YZ wrote the manuscript.

## Competing interests

The authors declare no competing interests.

## Acknowledgements

We thank the UTHSA Next Generation Sequencing Facilities for services rendered for production of the Hi-C, scHi-C and scRNA-seq data. We would also like to thank Dr. Bing Ren at University of California at San Diego for sharing us with their human scHi-C data. We are grateful to Dr. Myles Brown of Center for Functional Cancer Epigenetics at Dana-Farber Cancer Institute for reading the manuscript and providing suggestive comments. This project was partially supported by grants from NIH R01GM114142 and U54CA217297.

## Notes

### Competing Interest Statement

The authors have declared no competing interest.

## References

1. Dekker J, Rippe K, Dekker M, Kleckner N. Capturing chromosome conformation. Science. Feb 15;295(5558):1306-11 (2002).

2. Simonis M, Klous P, Splinter E, Moshkin Y, Willemsen R, de Wit E, van Steensel B, de Laat W. Nuclear organization of active and inactive chromatin domains uncovered by chromosome conformation capture-on-chip (4C). Nat Genet. Nov;38(11):1348–54 (2006).

3. Dostie J, Richmond TA, Arnaout RA, Selzer RR, Lee WL, Honan TA, Rubio ED, Krumm A, Lamb J, Nusbaum C, Green RD, Dekker J. Chromosome Conformation Capture Carbon Copy (5C): a massively parallel solution for mapping interactions between genomic elements. Genome Res. Oct;16(10):1299–309 (2006).

4. Fullwood MJ, Liu MH, Pan YF, Liu J, Xu H, Mohamed YB, Orlov YL, Velkov S, Ho A, Mei PH, Chew EG, Huang PY, Welboren WJ, Han Y, Ooi HS, Ariyaratne PN, Vega VB, Luo Y, Tan PY, Choy PY, Wansa KD, Zhao B, Lim KS, Leow SC, Yow JS, Joseph R, Li H, Desai KV, Thomsen JS, Lee YK, Karuturi RK, Herve T, Bourque G, Stunnenberg HG, Ruan X, Cacheux-Rataboul V, Sung WK, Liu ET, Wei CL, Cheung E, Ruan Y. An oestrogen-receptor-alpha-bound human chromatin interactome. Nature. Nov 5;462(7269):58-64 (2009).

5. Lieberman-Aiden E, van Berkum NL, Williams L, Imakaev M, Ragoczy T, Telling A, Amit I, Lajoie BR, Sabo PJ, Dorschner MO, Sandstrom R, Bernstein B, Bender MA, Groudine M, Gnirke A, Stamatoyannopoulos J, Mirny LA, Lander ES, Dekker J. Comprehensive mapping of long-range interactions reveals folding principles of the human genome. Science. Oct 9;326(5950):289-93 (2009).

6. Kalhor R, Tjong H, Jayathilaka N, Alber F, Chen L. Genome architectures revealed by tethered chromosome conformation capture and population-based modeling. Nat Biotechnol. Dec 25;30(1):90–8 (2011).

7. Rao SS, Huntley MH, Durand NC, Stamenova EK, Bochkov ID, Robinson JT, Sanborn AL, Machol I, Omer AD, Lander ES, Aiden EL. A 3D map of the human genome at kilobase resolution reveals principles of chromatin looping. Cell. Dec 18;159(7):1665–80 (2014).

8. Imakaev M, Fudenberg G, McCord RP, Naumova N, Goloborodko A, Lajoie BR, Dekker J, Mirny LA. Iterative correction of Hi-C data reveals hallmarks of chromosome organization. Nat Methods. Oct;9(10):999–1003 (2012).

9. Dixon JR, Selvaraj S, Yue F, Kim A, Li Y, Shen Y, Hu M, Liu JS, Ren B. Topological domains in mammalian genomes identified by analysis of chromatin interactions. Nature. Apr 11;485(7398):376-80 (2012).

10. Servant N, Lajoie BR, Nora EP, Giorgetti L, Chen CJ, Heard E, Dekker J, Barillot E. HiTC: exploration of high-throughput ’C’ experiments. Bioinformatics. Nov 1;28(21):2843–4 (2012).

11. Zheng X, Zheng Y. CscoreTool: fast Hi-C compartment analysis at high resolution. Bioinformatics. May 1;34(9):1568–1570 (2018).

12. Crane E, Bian Q, McCord RP, Lajoie BR, Wheeler BS, Ralston EJ, Uzawa S, Dekker J, Meyer BJ. Condensin-driven remodelling of X chromosome topology during dosage compensation. Nature. Jul 9;523(7559):240-4 (2015).

13. Shin H, Shi Y, Dai C, Tjong H, Gong K, Alber F, Zhou XJ. TopDom: an efficient and deterministic method for identifying topological domains in genomes. Nucleic Acids Res. Apr 20;44(7):e70 (2016).

14. Ay F, Bailey TL, Noble WS. Statistical confidence estimation for Hi-C data reveals regulatory chromatin contacts. Genome Res. Jun;24(6):999–1011 (2014).

15. Zhou Y, Cheng X, Yang Y, Li T, Li J, Huang TH, Wang J, Lin S, Jin VX. Modeling and analysis of Hi-C data by HiSIF identifies characteristic promoter-distal loops. Genome Med. Aug 12;12(1):69 (2020).

16. Zhang Y, An L, Xu J, Zhang B, Zheng WJ, Hu M, Tang J, Yue F. Enhancing Hi-C data resolution with deep convolutional neural network HiCPlus. Nat Commun. Feb 21;9(1):750 (2018).

17. Liu Q, Lv H, Jiang R. hicGAN infers super resolution Hi-C data with generative adversarial networks. Bioinformatics. Jul 15;35(14):i99–i107 (2019).

18. Durand NC, Robinson JT, Shamim MS, Machol I, Mesirov JP, Lander ES, Aiden EL. Juicebox Provides a Visualization System for Hi-C Contact Maps with Unlimited Zoom. Cell Syst. Jul;3(1):99–101 (2016).

19. Li D, Hsu S, Purushotham D, Sears RL, Wang T. WashU Epigenome Browser update 2019. Nucleic Acids Res. Jul 2;47(W1):W158–W165 (2019).

20. Wang Y, Song F, Zhang B, Zhang L, Xu J, Kuang D, Li D, Choudhary MNK, Li Y, Hu M, Hardison R, Wang T, Yue F. The 3D Genome Browser: a web-based browser for visualizing 3D genome organization and long-range chromatin interactions. Genome Biol. Oct 4;19(1):151 (2018).

21. Akdemir KC, Chin L. HiCPlotter integrates genomic data with interaction matrices. Genome Biol. Sep 21;16(1):198 (2015).

22. Nagano T, Lubling Y, Stevens TJ, Schoenfelder S, Yaffe E, Dean W, Laue ED, Tanay A, Fraser P. Single-cell Hi-C reveals cell-to-cell variability in chromosome structure. Nature. Oct 3;502(7469):59-64 (2013).

23. Ramani V, Deng X, Qiu R, Gunderson KL, Steemers FJ, Disteche CM, Noble WS, Duan Z, Shendure J. Massively multiplex single-cell Hi-C. Nat Methods. Mar;14(3):263–266 (2017).

24. Stevens TJ, Lando D, Basu S, Atkinson LP, Cao Y, Lee SF, Leeb M, Wohlfahrt KJ, Boucher W, O’Shaughnessy-Kirwan A, Cramard J, Faure AJ, Ralser M, Blanco E, Morey L, Sansó M, Palayret MGS, Lehner B, Di Croce L, Wutz A, Hendrich B, Klenerman D, Laue ED. 3D structures of individual mammalian genomes studied by single-cell Hi-C. *Nature*. Apr 6;544(7648):59-64 (2017).

25. Li G, Liu Y, Zhang Y, Kubo N, Yu M, Fang R, Kellis M, Ren B. Joint profiling of DNA methylation and chromatin architecture in single cells. Nat Methods. Oct;16(10):991–993 (2019).

26. Nagano T, Lubling Y, Várnai C, Dudley C, Leung W, Baran Y, Mendelson Cohen N, Wingett S, Fraser P, Tanay A. Cell-cycle dynamics of chromosomal organization at single-cell resolution. Nature. Jul 5;547(7661):61-67 (2017).

27. Flyamer IM, Gassler J, Imakaev M, Brandão HB, Ulianov SV, Abdennur N, Razin SV, Mirny LA, Tachibana-Konwalski K. Single-nucleus Hi-C reveals unique chromatin reorganization at oocyte-to-zygote transition. Nature. Apr 6;544(7648):110-114 (2017).

28. Gassler J, Brandão HB, Imakaev M, Flyamer IM, Ladstätter S, Bickmore WA, Peters JM, Mirny LA, Tachibana K. A mechanism of cohesin-dependent loop extrusion organizes zygotic genome architecture. EMBO J. Dec 15;36(24):3600–3618 (2017).

29. Bonora G, Ramani V, Singh R, Fang H, Jackson DL, Srivatsan S, Qiu R, Lee C, Trapnell C, Shendure J, Duan Z, Deng X, Noble WS, Disteche CM. Single-cell landscape of nuclear configuration and gene expression during stem cell differentiation and X inactivation. Genome Biol. Sep 27;22(1):279 (2021).

30. Allahyar A, Vermeulen C, Bouwman BAM, Krijger PHL, Verstegen MJAM, Geeven G, van Kranenburg M, Pieterse M, Straver R, Haarhuis JHI, Jalink K, Teunissen H, Renkens IJ, Kloosterman WP, Rowland BD, de Wit E, de Ridder J, de Laat W. Enhancer hubs and loop collisions identified from single-allele topologies. Nat Genet. Aug;50(8):1151–1160 (2018).

31. Oudelaar AM, Davies JOJ, Hanssen LLP, Telenius JM, Schwessinger R, Liu Y, Brown JM, Downes DJ, Chiariello AM, Bianco S, Nicodemi M, Buckle VJ, Dekker J, Higgs DR, Hughes JR. Single-allele chromatin interactions identify regulatory hubs in dynamic compartmentalized domains. Nat Genet. Dec;50(12):1744–1751 (2018).

32. Rosenthal M, Bryner D, Huffer F, Evans S, Srivastava A, Neretti N. Bayesian Estimation of Three-Dimensional Chromosomal Structure from Single-Cell Hi-C Data. J Comput Biol. Nov;26(11):1191–1202 (2019).

33. Zhu H, Wang Z. SCL: a lattice-based approach to infer 3D chromosome structures from single-cell Hi-C data. Bioinformatics. Oct 15;35(20):3981–3988 (2019).

34. Meng L, Wang C, Shi Y, Luo Q. Si-C is a method for inferring super-resolution intact genome structure from single-cell Hi-C data. Nat Commun. Jul 16;12(1):4369 (2021).

35. Liu J, Lin D, Yardimci GG, Noble WS. Unsupervised embedding of single-cell Hi-C data. Bioinformatics. Jul 1;34(13):i96–i104 (2018).

36. Zhou J, Ma J, Chen Y, Cheng C, Bao B, Peng J, Sejnowski TJ, Dixon JR, Ecker JR. Robust single-cell Hi-C clustering by convolution-and random-walk-based imputation. Proc Natl Acad Sci USA. Jul 9;116(28):14011–14018 (2019).

37. Zhang R, Zhou T, Ma J. Multiscale and integrative single-cell Hi-C analysis with Higashi. Nat Biotechnol. Oct 11. doi: 10.1038/s41587-021-01034-y. (2021)

38. Li X, Zeng G, Li A, Zhang Z. DeTOKI identifies and characterizes the dynamics of chromatin TAD-like domains in a single cell. Genome Biol. Jul 27;22(1):217<otherinfo> (2021)</otherinfo>.

39. Wu H, Wu Y, Jiang Y, Zhou B, Zhou H, Chen Z, Xiong Y, Liu Q, Zhang H. scHiCStackL: a stacking ensemble learning-based method for single-cell Hi-C classification using cell embedding. Brief Bioinform. Sep 22:bbab396 (2021).

40. Yu M, Abnousi A, Zhang Y, Li G, Lee L, Chen Z, Fang R, Lagler TM, Yang Y, Wen J, Sun Q, Li Y, Ren B, Hu M. SnapHiC: a computational pipeline to identify chromatin loops from single-cell Hi-C data. Nat Methods. Sep;18(9):1056–1059 (2021).

41. Li X, Feng F, Pu H, Leung WY, Liu J. scHiCTools: A computational toolbox for analyzing single-cell Hi-C data. PLoS Comput Biol. May 18;17(5):e1008978 (2021).

42. Niveditha D, Sharma H, Sahu A, Majumder S, Chowdhury R, Chowdhury S. Drug tolerant cells: an emerging target with unique transcriptomic features. Cancer Inform. Oct 10;18:1176935119881633 (2019).

43. Xue Y, Martelotto L, Baslan T, Vides A, Solomon M, Mai TT, Chaudhary N, Riely GJ, Li BT, Scott K, Cechhi F, Stierner U, Chadalavada K, de Stanchina E, Schwartz S, Hembrough T, Nanjangud G, Berger MF, Nilsson J, Lowe SW, Reis-Filho JS, Rosen N, Lito P. An approach to suppress the evolution of resistance in BRAF(V600E)-mutant cancer. Nat Med. Aug;23(8):929–937 (2017).

44. Shaffer SM, Dunagin MC, Torborg SR, Torre EA, Emert B, Krepler C, Beqiri M, Sproesser K, Brafford PA, Xiao M, Eggan E, Anastopoulos IN, Vargas-Garcia CA, Singh A, Nathanson KL, Herlyn M, Raj A. Rare cell variability and drug-induced reprogramming as a mode of cancer drug resistance. Nature. Jun 15;546(7658):431-435 (2017).

45. Oren Y, Tsabar M, Cuoco MS, Amir-Zilberstein L, Cabanos HF, Hütter JC, Hu B, Thakore PI, Tabaka M, Fulco CP, Colgan W, Cuevas BM, Hurvitz SA, Slamon DJ, Deik A, Pierce KA, Clish C, Hata AN, Zaganjor E, Lahav G, Politi K, Brugge JS, Regev A. Cycling cancer persister cells arise from lineages with distinct programs. Nature. Aug;596(7873):576-582 (2021).

46. Zhou Y, Gerrard DL, Wang J, Li T, Yang Y, Fritz AJ, Rajendran M, Fu X, Stein G, Schiff R, Lin S, Frietze S, Jin VX. Temporal dynamic reorganization of 3D chromatin architecture in hormone-induced breast cancer and endocrine resistance. Nat Commun. Apr 3;10(1):1522 (2019).

47. Kilker RL, Planas-Silva MD. Cyclin D1 is necessary for tamoxifen-induced cell cycle progression in human breast cancer cells. Cancer Res. Dec 1;66(23):11478–84 (2006).

48. Ferraiuolo RM, Tubman J, Sinha I, Hamm C, Porter LA. The cyclin-like protein, SPY1, regulates the ERα and ERK1/2 pathways promoting tamoxifen resistance. Oncotarget. Apr 4;8(14):23337–23352 (2017).

49. Løkkegaard S, Elias D, Alves CL, Bennetzen MV, Lænkholm AV, Bak M, Gjerstorff MF, Johansen LE, Vever H, Bjerre C, Kirkegaard T, Nordenskjöld B, Fornander T, Stål O, Lindström LS, Esserman LJ, Lykkesfeldt AE, Andersen JS, Leth-Larsen R, Ditzel HJ. MCM3 upregulation confers endocrine resistance in breast cancer and is a predictive marker of diminished tamoxifen benefit. NPJ Breast Cancer. Jan 4;7(1):2 (2021).

50. Gao A, Sun T, Ma G, Cao J, Hu Q, Chen L, Wang Y, Wang Q, Sun J, Wu R, Wu Q, Zhou J, Liu L, Hu J, Dong JT, Zhu Z. LEM4 confers tamoxifen resistance to breast cancer cells by activating cyclin D-CDK4/6-Rb and ERα pathway. Nat Commun. Oct 9;9(1):4180 (2018).

51. Yu D, Shi L, Bu Y, Li W. Cell Division Cycle Associated 8 Is a Key Regulator of Tamoxifen Resistance in Breast Cancer. J Breast Cancer. Jun 7;22(2):237–247 (2019).

52. Bi M, Zhang Z, Jiang YZ, Xue P, Wang H, Lai Z, Fu X, De Angelis C, Gong Y, Gao Z, Ruan J, Jin VX, Marangoni E, Montaudon E, Glass CK, Li W, Huang TH, Shao ZM, Schiff R, Chen L, Liu Z. Enhancer reprogramming driven by high-order assemblies of transcription factors promotes phenotypic plasticity and breast cancer endocrine resistance. Nat Cell Biol. Jun;22(6):701–715 (2020).

53. Li J, Fang K, Choppavarapu L, Yang K, Yang Y, Wang J, Cao R, Jatoi I, Jin VX. Hi-C profiling of cancer spheroids identifies 3D-growth-specific chromatin interactions in breast cancer endocrine resistance. Clin Epigenetics. Sep 17;13(1):175 (2021).

54. Yang Y, Choppavarapu L, Fang K, Naeini AS, Nosirov B, Li J, Yang K, He Z, Zhou Y, Schiff R, Li R, Hu Y, Wang J, Jin VX. The 3D genomic landscape of differential response to EGFR/HER2 inhibition in endocrine-resistant breast cancer cells. Biochim Biophys Acta Gene Regul Mech. Nov;1863(11):194631 (2020).

55. Montero-Conde C, Leandro-Garcia LJ, Chen X, Oler G, Ruiz-Llorente S, Ryder M, Landa I, Sanchez-Vega F, La K, Ghossein RA, Bajorin DF, Knauf JA, Riordan JD, Dupuy AJ, Fagin JA. Transposon mutagenesis identifies chromatin modifiers cooperating with Ras in thyroid tumorigenesis and detects ATXN7 as a cancer gene. Proc Natl Acad Sci USA. Jun 20;114(25):E4951–E4960 (2017).

56. Atanassov BS, Mohan RD, Lan X, Kuang X, Lu Y, Lin K, McIvor E, Li W, Zhang Y, Florens L, Byrum SD, Mackintosh SG, Calhoun-Davis T, Koutelou E, Wang L, Tang DG, Tackett AJ, Washburn MP, Workman JL, Dent SY. ATXN7L3 and ENY2 Coordinate Activity of Multiple H2B Deubiquitinases Important for Cellular Proliferation and Tumor Growth. Mol Cell. May 19;62(4):558–71 (2016).

57. Okuno K, Akiyama Y, Shimada S, Nakagawa M, Tanioka T, Inokuchi M, Yamaoka S, Kojima K, Tanaka S. Asymmetric dimethylation at histone H3 arginine 2 by PRMT6 in gastric cancer progression. Carcinogenesis. Mar 12;40(1):15–26 (2019).

58. Jiang N, Li QL, Pan W, Li J, Zhang MF, Cao T, Su SG, Shen H. PRMT6 promotes endometrial cancer via AKT/mTOR signaling and indicates poor prognosis. Int J Biochem Cell Biol. Mar;120:105681 (2020).

59. Avasarala S, Wu PY, Khan SQ, Yanlin S, Van Scoyk M, Bao J, Di Lorenzo A, David O, Bedford MT, Gupta V, Winn RA, Bikkavilli RK. PRMT6 Promotes Lung Tumor Progression via the Alternate Activation of Tumor-Associated Macrophages. Mol Cancer Res. Jan;18(1):166–178 (2020).

60. Gallo M, Coutinho FJ, Vanner RJ, Gayden T, Mack SC, Murison A, Remke M, Li R, Takayama N, Desai K, Lee L, Lan X, Park NI, Barsyte-Lovejoy D, Smil D, Sturm D, Kushida MM, Head R, Cusimano MD, Bernstein M, Clarke ID, Dick JE, Pfister SM, Rich JN, Arrowsmith CH, Taylor MD, Jabado N, Bazett-Jones DP, Lupien M, Dirks PB. MLL5 Orchestrates a Cancer Self-Renewal State by Repressing the Histone Variant H3.3 and Globally Reorganizing Chromatin. Cancer Cell. Dec 14;28(6):715–729 (2015).

61. Takawa M, Cho HS, Hayami S, Toyokawa G, Kogure M, Yamane Y, Iwai Y, Maejima K, Ueda K, Masuda A, Dohmae N, Field HI, Tsunoda T, Kobayashi T, Akasu T, Sugiyama M, Ohnuma S, Atomi Y, Ponder BA, Nakamura Y, Hamamoto R. Histone lysine methyltransferase SETD8 promotes carcinogenesis by deregulating PCNA expression. Cancer Res. Jul 1;72(13):3217–27 (2012).

62. Chen YY, Wang WH, Che L, Lan Y, Zhang LY, Zhan DL, Huang ZY, Lin ZN, Lin YC. BNIP3L-Dependent Mitophagy Promotes HBx-Induced Cancer Stemness of Hepatocellular Carcinoma Cells via Glycolysis Metabolism Reprogramming. Cancers (Basel). Mar 11;12(3):655 (2020).

63. Wagener N, Bulkescher J, Macher-Goeppinger S, Karapanagiotou-Schenkel I, Hatiboglu G, Abdel-Rahim M, Abol-Enein H, Ghoneim MA, Bastian PJ, Müller SC, Haferkamp A, Hohenfellner M, Hoppe-Seyler F, Hoppe-Seyler K. Endogenous BTG2 expression stimulates migration of bladder cancer cells and correlates with poor clinical prognosis for bladder cancer patients. Br J Cancer. Mar 5;108(4):973–82 (2013).

64. Ackermann T, Hartleben G, Müller C, Mastrobuoni G, Groth M, Sterken BA, Zaini MA, Youssef SA, Zuidhof HR, Krauss SR, Kortman G, de Haan G, de Bruin A, Wang ZQ, Platzer M, Kempa S, Calkhoven CF. C/EBPβ-LIP induces cancer-type metabolic reprogramming by regulating the let-7/LIN28B circuit in mice. Commun Biol. Jun 14;2:208 (2019).

65. Ikeda K, Horie-Inoue K, Suzuki T, Hobo R, Nakasato N, Takeda S, Inoue S. Mitochondrial supercomplex assembly promotes breast and endometrial tumorigenesis by metabolic alterations and enhanced hypoxia tolerance. Nat Commun. Sep 11;10(1):4108 (2019).

66. Banerjee S, Wei T, Wang J, Lee JJ, Gutierrez HL, Chapman O, Wiley SE, Mayfield JE, Tandon V, Juarez EF, Chavez L, Liang R, Sah RL, Costello C, Mesirov JP, de la Vega L, Cooper KL, Dixon JE, Xiao J, Lei X. Inhibition of dual-specificity tyrosine phosphorylation-regulated kinase 2 perturbs 26S proteasome-addicted neoplastic progression. Proc Natl Acad Sci USA. Dec 3;116(49):24881–24891 (2019).

67. Rasool RU, Natesan R, Deng Q, Aras S, Lal P, Sander Effron S, Mitchell-Velasquez E, Posimo JM, Carskadon S, Baca SC, Pomerantz MM, Siddiqui J, Schwartz LE, Lee DJ, Palanisamy N, Narla G, Den RB, Freedman ML, Brady DC, Asangani IA. CDK7 Inhibition Suppresses Castration-Resistant Prostate Cancer through MED1 Inactivation. Cancer Discov. Nov;9(11):1538–1555 (2019).

68. Xiang Y, Ye Y, Lou Y, Yang Y, Cai C, Zhang Z, Mills T, Chen NY, Kim Y, Muge Ozguc F, Diao L, Karmouty-Quintana H, Xia Y, Kellems RE, Chen Z, Blackburn MR, Yoo SH, Shyu AB, Mills GB, Han L. Comprehensive Characterization of Alternative Polyadenylation in Human Cancer. J Natl Cancer Inst. Apr 1;110(4):379–389 (2018).

69. Canevari RA, Marchi FA, Domingues MA, de Andrade VP, Caldeira JR, Verjovski-Almeida S, Rogatto SR, Reis EM. Identification of novel biomarkers associated with poor patient outcomes in invasive breast carcinoma. Tumour Biol. Oct;37(10):13855–13870 (2016).

70. Hao Y, Hao S, Andersen-Nissen E, Mauck WM 3rd, Zheng S, Butler A, Lee MJ, Wilk AJ, Darby C, Zager M, Hoffman P, Stoeckius M, Papalexi E, Mimitou EP, Jain J, Srivastava A, Stuart T, Fleming LM, Yeung B, Rogers AJ, McElrath JM, Blish CA, Gottardo R, Smibert P, Satija R. Integrated analysis of multimodal single-cell data. Cell. Jun 24;184(13):3573–3587.e29 (2021).

71. Servant N, Varoquaux N, Lajoie BR, Viara E, Chen CJ, Vert JP, Heard E, Dekker J, Barillot E. HiC-Pro: an optimized and flexible pipeline for Hi-C data processing. Genome Biol. Dec 1;16:259 (2015).

72. Yang T, Zhang F, Yardımcı GG, Song F, Hardison RC, Noble WS, Yue F, Li Q. HiCRep: assessing the reproducibility of Hi-C data using a stratum-adjusted correlation coefficient. Genome Res. Nov;27(11):1939–1949 (2017).

73. Nora EP, Caccianini L, Fudenberg G, So K, Kameswaran V, Nagle A, Uebersohn A, Hajj B, Saux AL, Coulon A, Mirny LA, Pollard KS, Dahan M, Bruneau BG. Molecular basis of CTCF binding polarity in genome folding. Nat Commun. Nov 5;11(1):5612 (2020).

74. Whyte WA, Orlando DA, Hnisz D, Abraham BJ, Lin CY, Kagey MH, Rahl PB, Lee TI, Young RA. Master transcription factors and mediator establish super-enhancers at key cell identity genes. Cell. Apr 11;153(2):307–19 (2013).

75. Massarweh S, Osborne CK, Creighton CJ, Qin L, Tsimelzon A, Huang S, Weiss H, Rimawi M, Schiff R. Tamoxifen resistance in breast tumors is driven by growth factor receptor signaling with repression of classic estrogen receptor genomic function. Cancer Res. Feb 1;68(3):826–33 (2008).

76. Feng Q, Zhang Z, Shea MJ, Creighton CJ, Coarfa C, Hilsenbeck SG, Lanz R, He B, Wang L, Fu X, Nardone A, Song Y, Bradner J, Mitsiades N, Mitsiades CS, Osborne CK, Schiff R, O’Malley BW. An epigenomic approach to therapy for tamoxifen-resistant breast cancer. Cell Res. Jul;24(7):809–19 (2014).

77. Morrison G, Fu X, Shea M, Nanda S, Giuliano M, Wang T, Klinowska T, Osborne CK, Rimawi MF, Schiff R. Therapeutic potential of the dual EGFR/HER2 inhibitor AZD8931 in circumventing endocrine resistance. Breast Cancer Res Treat. Apr;144(2):263–72 (2014).

78. Kumar G, Garnova E, Reagin M, Vidali A. Improved multiple displacement amplification with phi29 DNA polymerase for genotyping of single human cells. Biotechniques. Jun;44(7):879–90 (2008).

79. Subramanian A, Tamayo P, Mootha VK, Mukherjee S, Ebert BL, Gillette MA, Paulovich A, Pomeroy SL, Golub TR, Lander ES, Mesirov JP. Gene set enrichment analysis: a knowledge-based approach for interpreting genome-wide expression profiles. Proc Natl Acad Sci USA. Oct 25;102(43):15545–50 (2005).

80. Sotiriou C, Wirapati P, Loi S, Harris A, Fox S, Smeds J, Nordgren H, Farmer P, Praz V, Haibe-Kains B, Desmedt C, Larsimont D, Cardoso F, Peterse H, Nuyten D, Buyse M, Van de Vijver MJ, Bergh J, Piccart M, Delorenzi M. Gene expression profiling in breast cancer: understanding the molecular basis of histologic grade to improve prognosis. J Natl Cancer Inst. Feb 15;98(4):262–72 (2006).

81. Loi S, Haibe-Kains B, Desmedt C, Wirapati P, Lallemand F, Tutt AM, Gillet C, Ellis P, Ryder K, Reid JF, Daidone MG, Pierotti MA, Berns EM, Jansen MP, Foekens JA, Delorenzi M, Bontempi G, Piccart MJ, Sotiriou C. Predicting prognosis using molecular profiling in estrogen receptor-positive breast cancer treated with tamoxifen. BMC Genomics. May 22;9:239 (2008).

82. Oliveira Junior AB, Contessoto VG, Mello MF, Onuchic JN. A Scalable Computational Approach for Simulating Complexes of Multiple Chromosomes. J Mol Biol. Mar 19;433(6):166700 (2021).

83. Pettersen EF, Goddard TD, Huang CC, Couch GS, Greenblatt DM, Meng EC, Ferrin TE. UCSF Chimera--a visualization system for exploratory research and analysis. J Comput Chem. Oct;25(13):1605–12 (2004).

84. Friedman CE, Nguyen Q, Lukowski SW, Helfer A, Chiu HS, Miklas J, Levy S, Suo S, Han JJ, Osteil P, Peng G, Jing N, Baillie GJ, Senabouth A, Christ AN, Bruxner TJ, Murry CE, Wong ES, Ding J, Wang Y, Hudson J, Ruohola-Baker H, Bar-Joseph Z, Tam PPL, Powell JE, Palpant NJ. Single-Cell Transcriptomic Analysis of Cardiac Differentiation from Human PSCs Reveals HOPX-Dependent Cardiomyocyte Maturation. Cell Stem Cell. Oct 4;23(4):586–598.e8 (2018).

85. Gonzalez-Pena V, Natarajan S, Xia Y, Klein D, Carter R, Pang Y, Shaner B, Annu K, Putnam D, Chen W, Connelly J, Pruett-Miller S, Chen X, Easton J, Gawad C. Accurate genomic variant detection in single cells with primary template-directed amplification. Proc Natl Acad Sci U S A. 2021 Jun 15;118(24).

86. Dong X, Zhang L, Hao X, Wang T, Vijg J. SCCNV: A Software Tool for Identifying Copy Number Variation From Single-Cell Whole-Genome Sequencing. Front Genet. 2020 Nov 16;11:505441.

87. Lee DS, Luo C, Zhou J, Chandran S, Rivkin A, Bartlett A, Nery JR, Fitzpatrick C, O’Connor C, Dixon JR, Ecker JR. Simultaneous profiling of 3D genome structure and DNA methylation in single human cells. Nat Methods. 2019 Oct;16(10):999–1006.

88. Darmanis S, Sloan SA, Zhang Y, Enge M, Caneda C, Shuer LM, Hayden Gephart MG, Barres BA, Quake SR. A survey of human brain transcriptome diversity at the single cell level. Proc Natl Acad Sci U S A. 2015 Jun 9;112(23):7285–90.

89. Kumegawa K, Takahashi Y, Saeki S, Yang L, Nakadai T, Osako T, Mori S, Noda T, Ohno S, Ueno T, Maruyama R. GRHL2 motif is associated with intratumor heterogeneity of cis-regulatory elements in luminal breast cancer. NPJ Breast Cancer. 2022 Jun 8;8(1):70.

