## Supplementary material for "Integration of scHi-C and scRNA-seq data defines distinct 3D-regulated and biological-context dependent cell subpopulations": Suppl. Figures

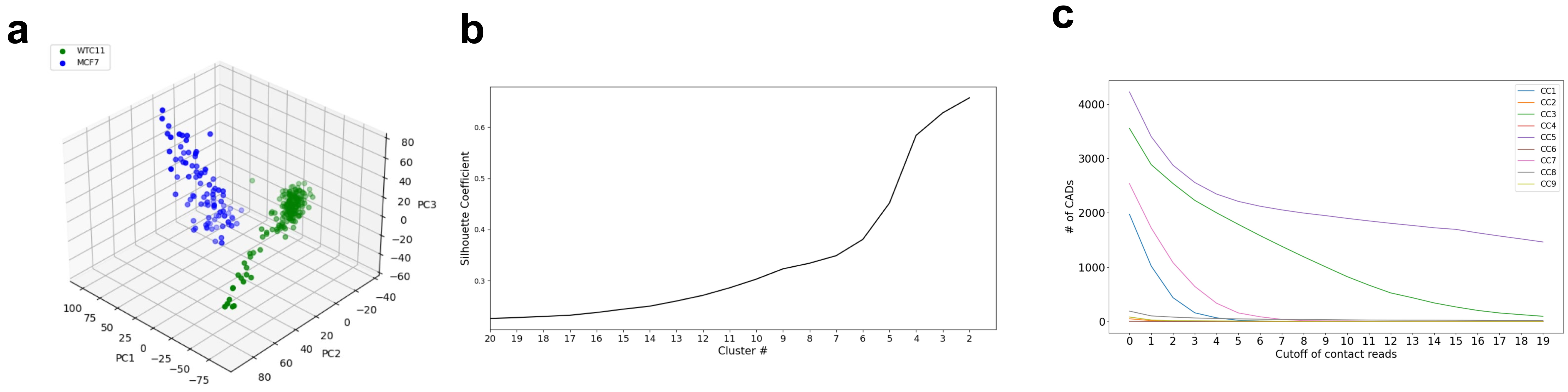


**d**

| scHi-C / scRNAseq | WTC11<br>CC1, CC3, CC5, CC7 | MCF7<br>CC2, CC4, CC6, CC8, CC9 |
|---|---|---|
| WTC11<br>DD1, DD2, DD4, DD5, DD7, DD8, DD9 | WMG1<br>622 genes | WMG3<br>127 genes |
| MCF7<br>DD3, DD6, DD10 | WMG2<br>1810 genes | WMG4<br>479 genes |

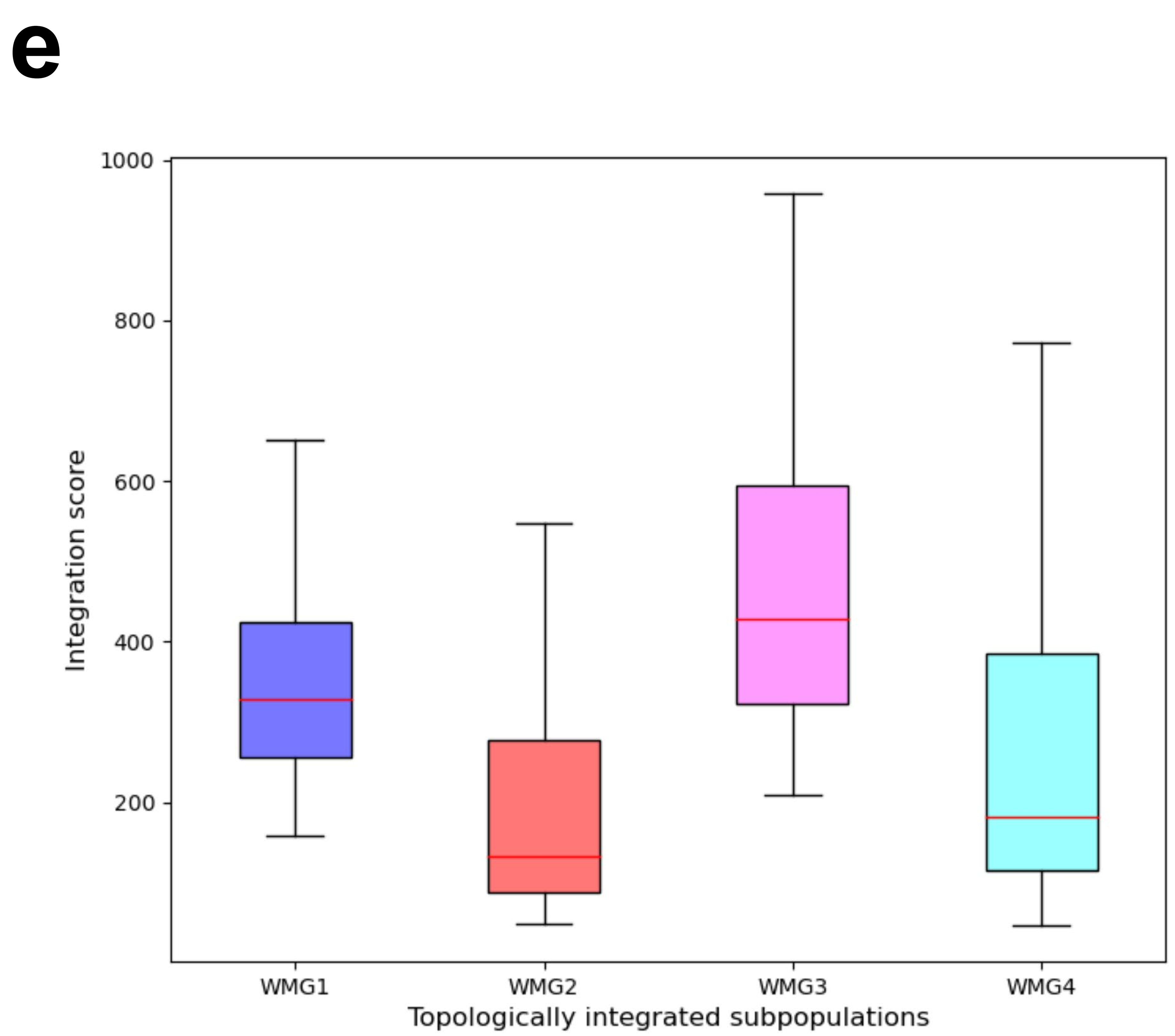

**Extended Data Fig. 1. Integration of scHi-C and scRNA-seq for WTC11 and MCF7 cells.**

**a**, 3D view of scHi-C data of WTC11 and MCF7. **b**, The Silhouette Coefficient of cluster numbers for scHi-C data. **c**, # of CADs in various cutoff of contact reads for each cluster. **d**, Four TISPs (WMG1-4) identified by MUDI after the integration of scHi-C and scRNA-seq. **e**, Integration scores of four TISPs (WMG1-4).

**a**

| scHi-C / scRNAseq | WTC11 with Yamanaka factors CC3, CC5 | WTC11 without Yamanaka factors CC1, CC7 | MCF7 with Yamanaka factors CC8 | MCF7 without Yamanaka factors CC2, CC4, CC6, CC9 |
|---|---|---|---|---|
| WTC11 with Yamanaka factors DD1 | YFG1 | YFG4 | YFG7 | YFG10 |
| WTC11 without Yamanaka factors DD2, DD4, DD5, DD7, DD8, DD9 | YFG2 | YFG5 | YFG8 | YFG11 |
| MCF7 without Yamanaka factors DD3, DD6, DD10 | YFG3 | YFG6 | YFG9 | YFG12 |

**b**

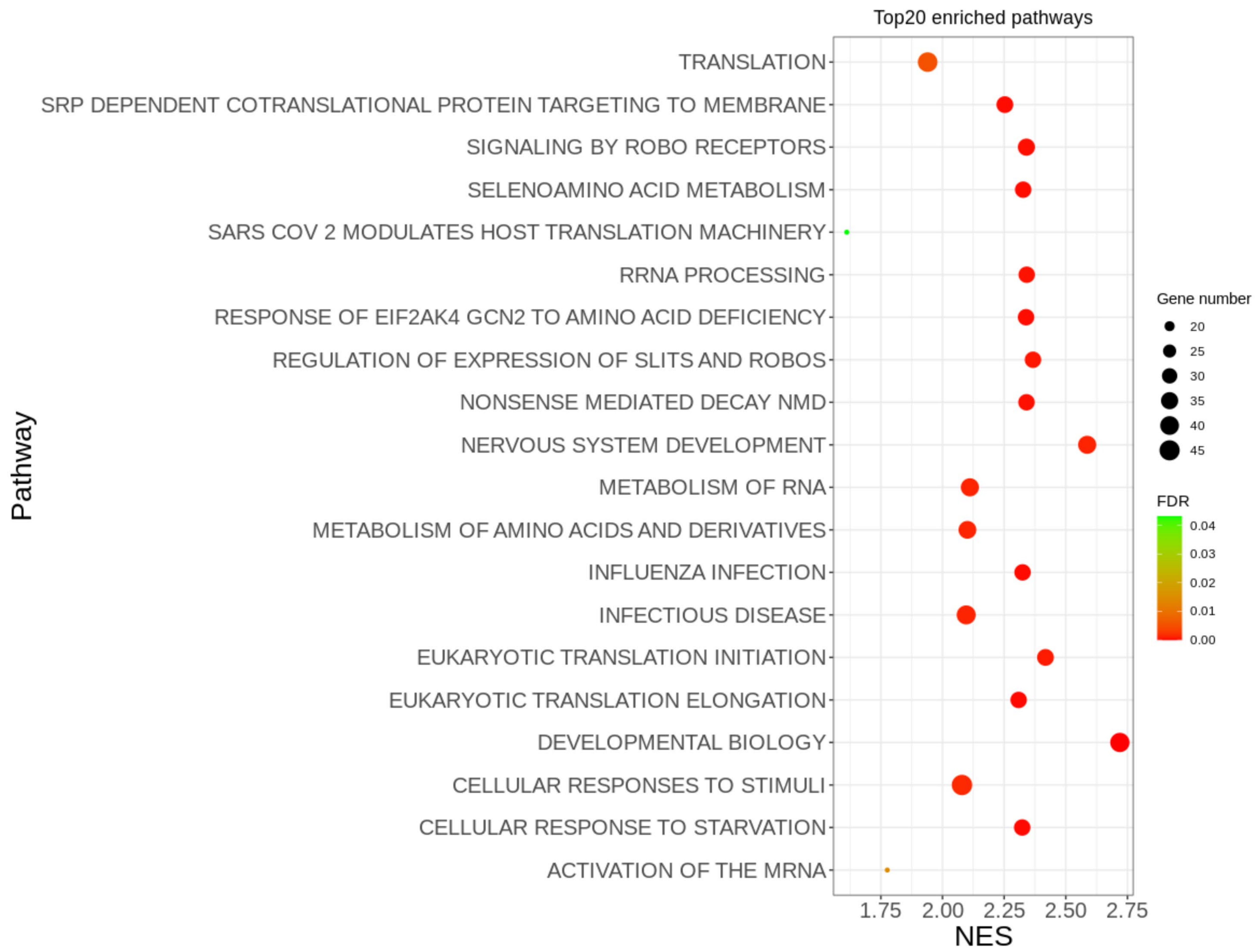


**c**

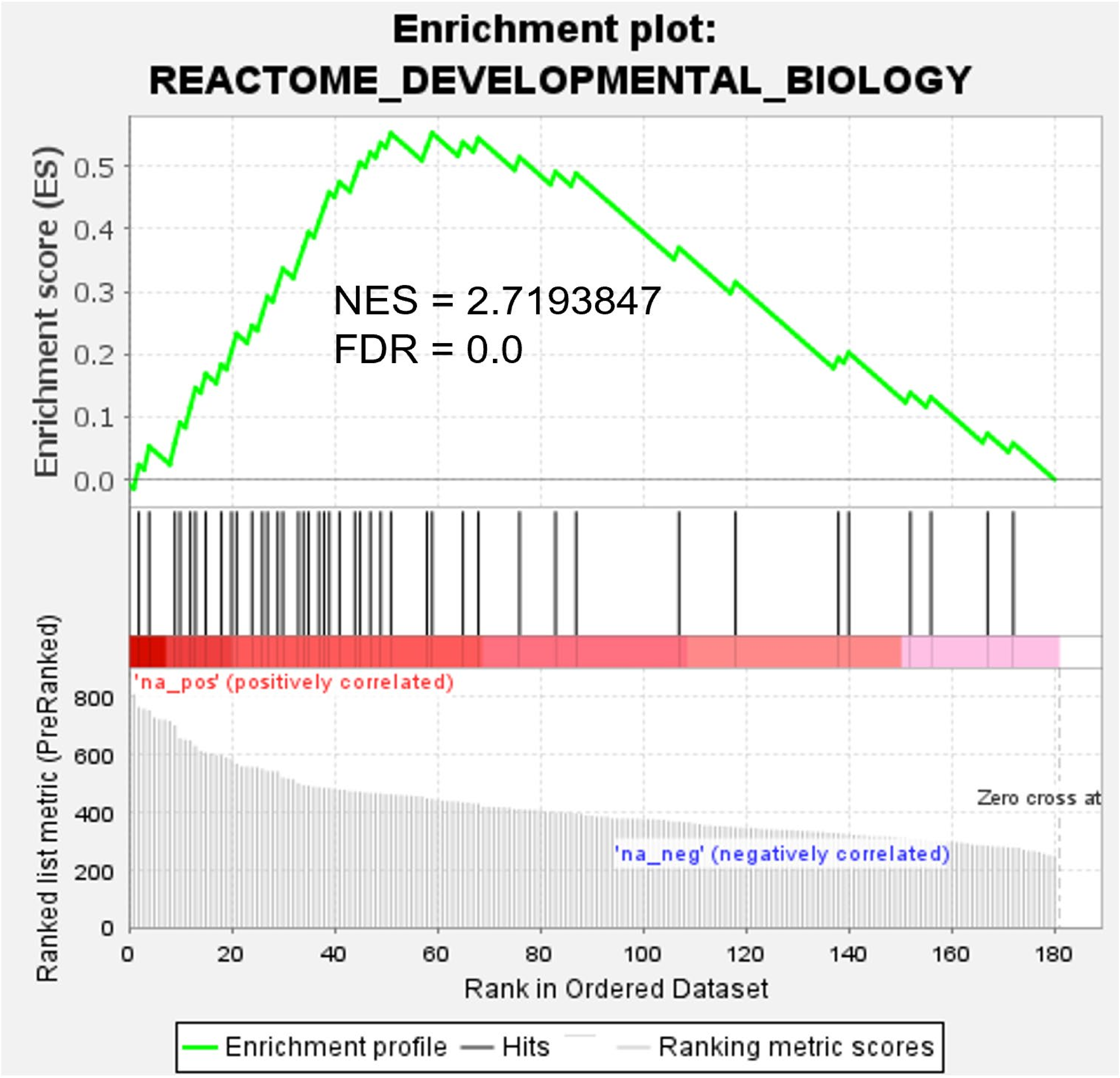

**Extended Data Fig. 2. 3D-regulated cell subpopulations based on Yamanaka factors.**

**a**, Twelve subpopulations (YFG1-12) identified by integration of scHi-C and scRNA-seq and Yamanaka factors. **b**, Enrichment of REACTOME signaling pathway for YFG1. **c**, YFG1 was majorly enriched to REACTOME Developmental Biology signaling pathway.

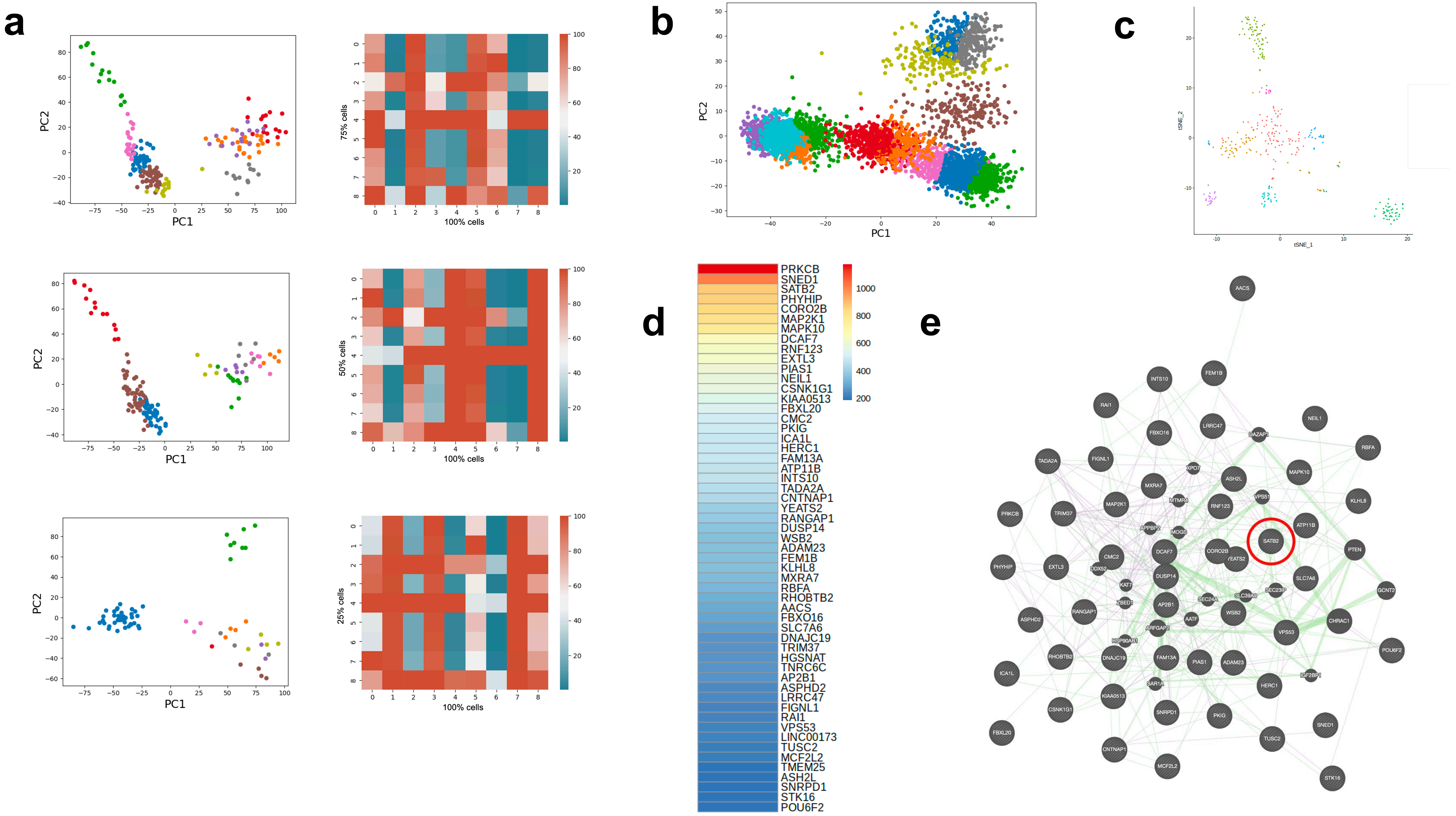

a
PC1
PC2
75% cells
50% cells
25% cells
100% cells
b
c
tSNE_1
tSNE_2
d
PRKCB
SNED1
SATB2
PHYHIP
CORO2B
MAP2K1
MAPK10
DCAF7
RNF123
EXTL3
PIAS1
NEIL1
CSNK1G1
KIAA0513
FBXL20
CMC2
PKIG
ICA1L
HERC1
FAM13A
ATP11B
INTS10
TADA2A
CNTNAP1
YEATS2
RANGAP1
DUSP14
WSB2
ADAM23
FEM1B
KLHL8
MXRA7
RBFA
RHOBTB2
AACS
FBXO16
SLC7A6
DNAJC19
TRIM37
HGSNAT
TNRC6C
AP2B1
ASPHD2
LRRC47
FIGNL1
RAI1
VPS53
LINC00173
TUSC2
MCF2L2
TMEM25
ASH2L
SNRPD1
STK16
POU6F2
1000
800
600
400
200
e

**Extended Data Fig. 3. A sub-sampling and applied algorithm on brain tissue.**

**a**, Sub-sampling of WTC11 and MCF7 cells to 75% cells (208 cells, upper panels), 50% cells (138 cells, middle panels) and 25% cells (69 cells, lower panels). Left panels were 2D view of scHi-C data, right panels were the overlapped CADs (%) in each cluster. **b**, 2D view of human cortex sn-m3C-seq data. **c**, 2D view of human cortex scRNA-seq data. **d**, Integration scores of genes in TISPs with cell markers of excitatory neurons. **e**, Interaction networks of genes in TISPs with cell markers of excitatory neurons. One of excitatory neuron cell marker SATB2 was highlighted with red circle.

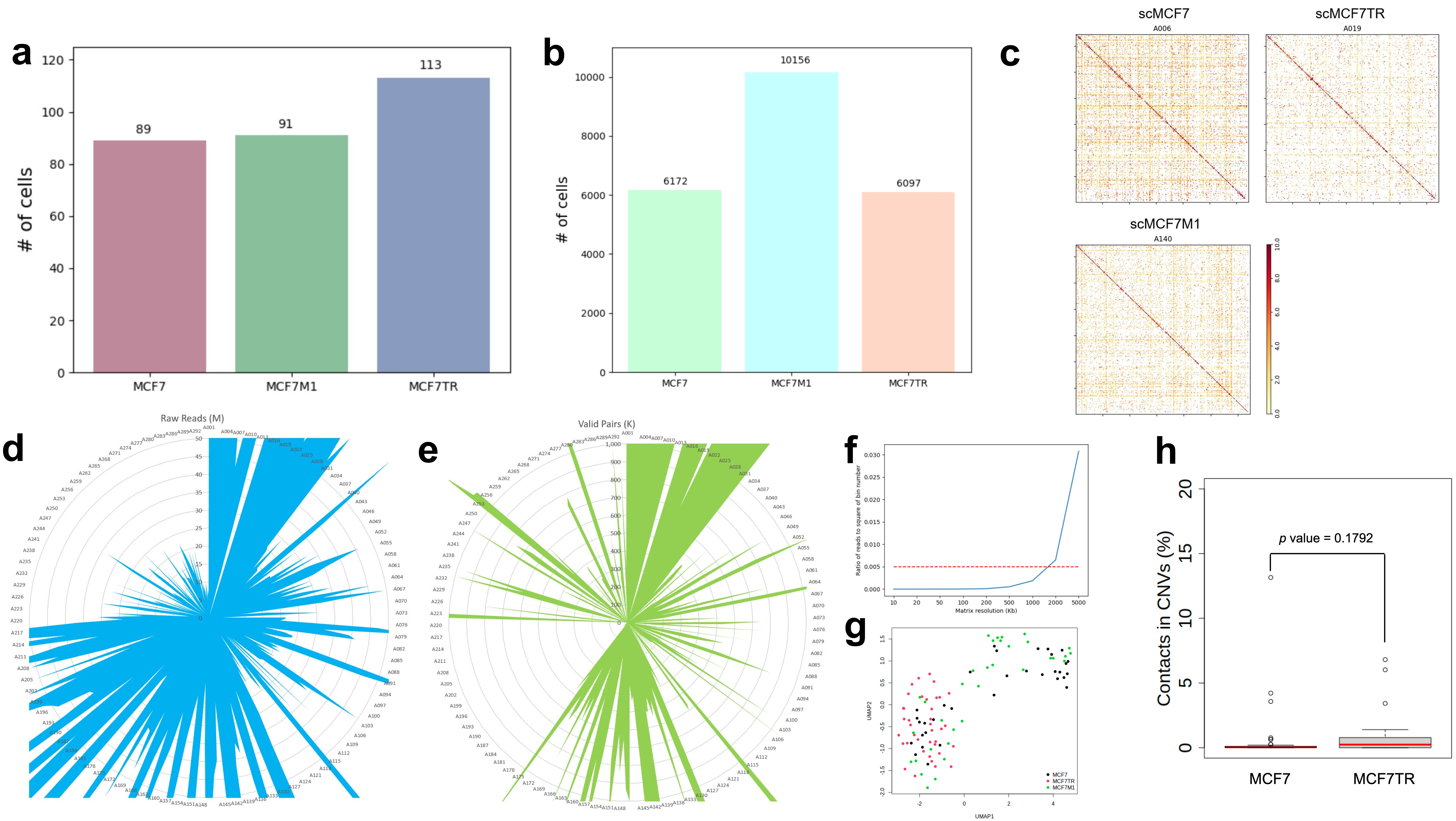

a
### of cells
89
91
113
MCF7
MCF7M1
MCF7TR
b
### of cells
6172
10156
6097
MCF7
MCF7M1
MCF7TR
c
scMCF7
A006
scMCF7TR
A019
scMCF7M1
A140
d
Raw Reads (M)
e
Valid Pairs (K)
f
Ratio of reads to square of bin number
Matrix resolution (Kb)
g
UMAP1
UMAP2
MCF7
MCF7TR
MCF7M1
h
Contacts in CNVs (%)
p value = 0.1792
MCF7
MCF7TR

**Extended Data Fig. 4. Summary of single-cell Hi-C data.**

**a**, Number of cells of scHi-C data. **b**, Number of cells of scRNA-seq data. **c**, Three examples of single cell contact maps: A006 – a MCF7 single cell, A140 – a MCF7M1 single cell, and A019 – a MCF7TR single cell. **d**, Raw reads of scHi-C data. Individual cells were labelled with the prefix of A and sequence number 001-293. **e**, Valid pairs of scHi-C data. **f**, Ratio of reads to square of bin number of scHi-C data in various matrix resolution. Red line was the cutoff of the sparsity, the indicated maximal resolution under the red line is 1Mb, which was used for the clustering of scHi-C data. **g**. 2D view of copy number variations (CNVs) of scDNA-seq data in MCF7, MCF7M1 and MCF7TR cells indicating there was no clear difference among cells. **h**. scHi-C contacts in the regions where 10% cells have CNVs. *p* value was determined by Wilcoxon rank-sum test.

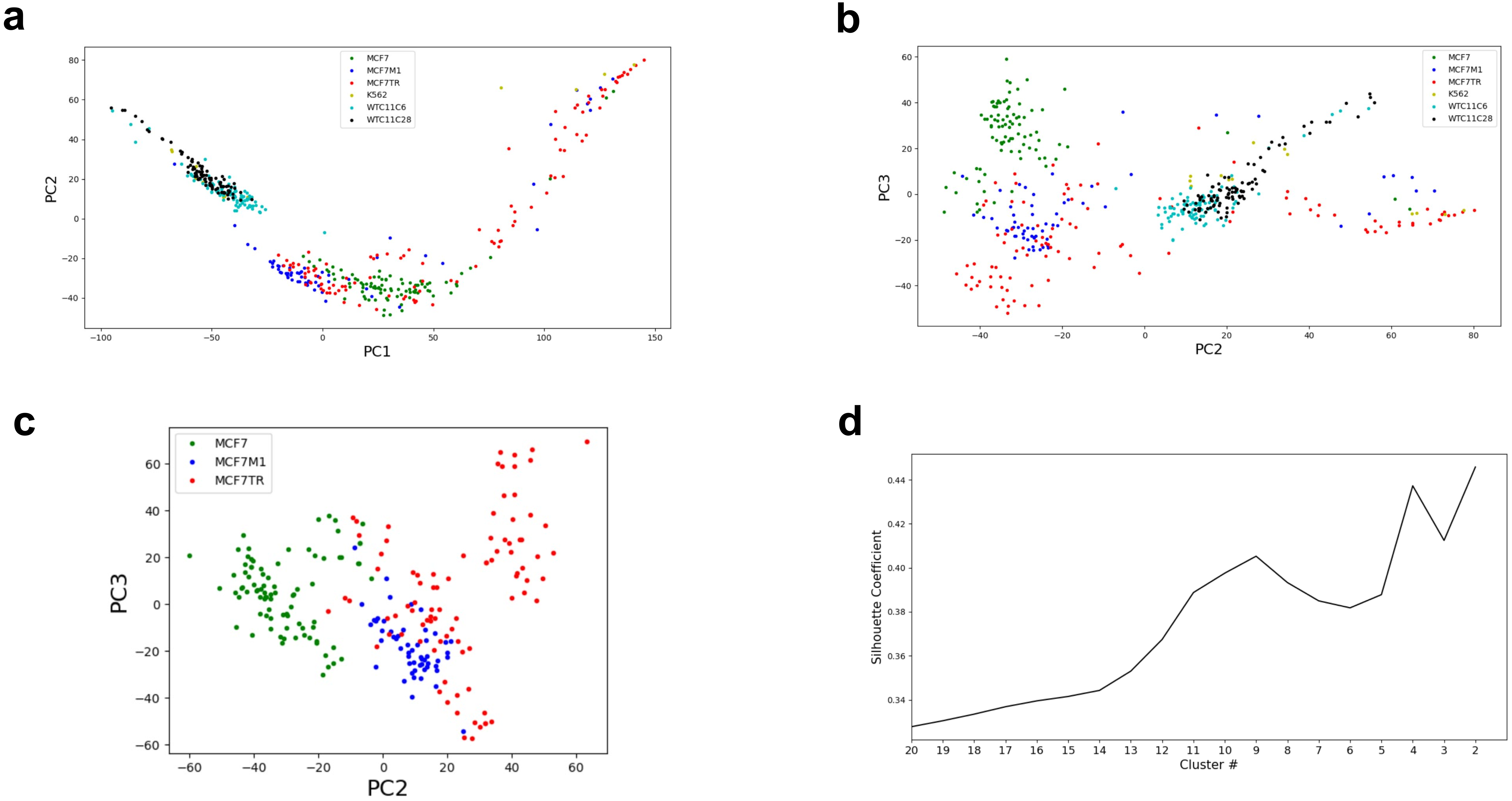
a
b
c
d
MCF7
MCF7M1
MCF7TR
K562
WTC11C6
WTC11C28
PC1
PC2
PC3
Silhouette Coefficient
Cluster #

**Extended Data Fig. 5. The view of scHi-C data from other dimensions.**

**a**, Comparing scHi-C data of this study with public human scHi-C data in the view of the first eigenvector (PC1) and the second eigenvector (PC2). **b**, Comparing scHi-C data of this study with public human scHi-C data in the view of the second eigenvector (PC2) and the third eigenvector (PC3). **c**, scHi-C data in the view of the second eigenvector (PC2) and the third eigenvector (PC3). **d**, The Silhouette Coefficient of cluster numbers.

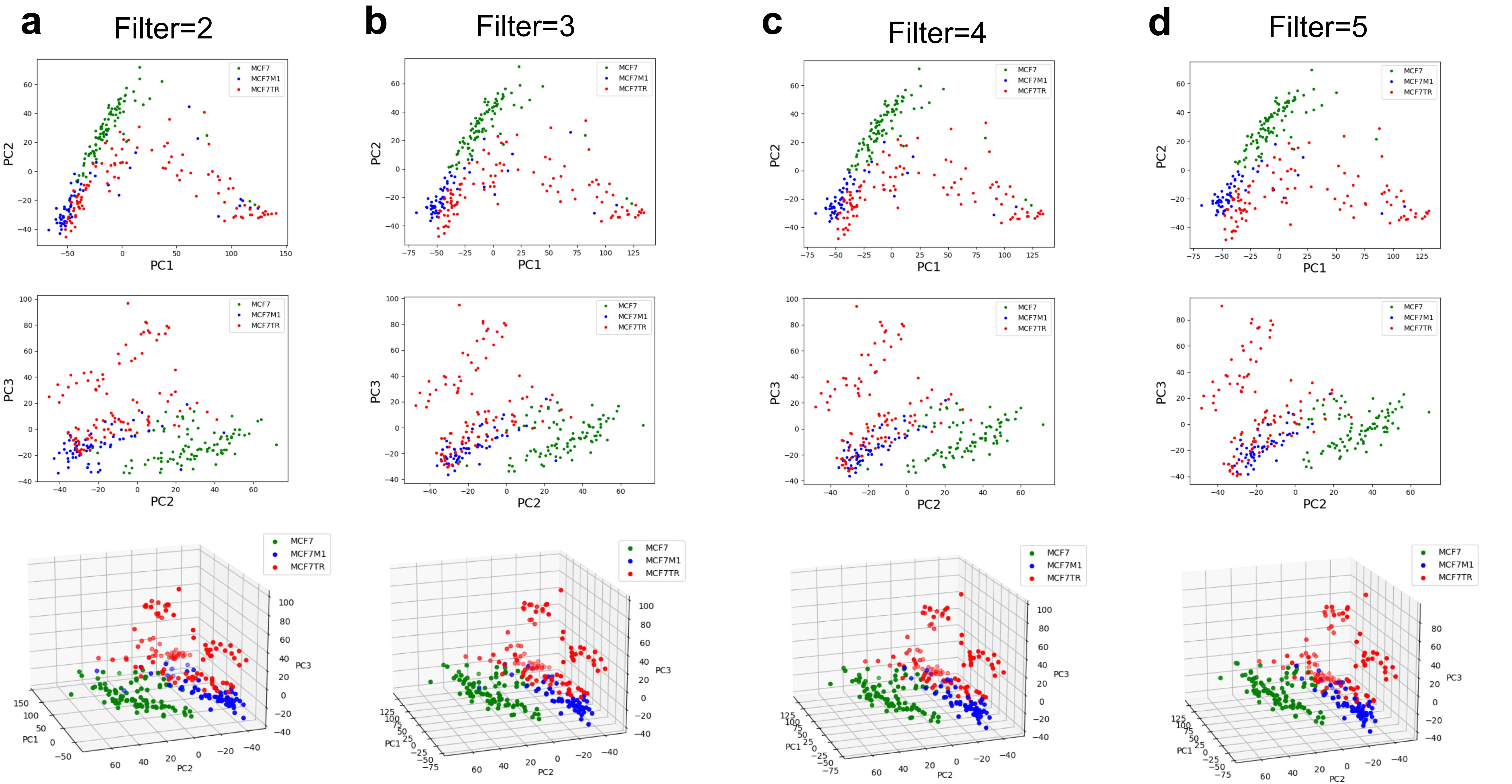

a
Filter=2
b
Filter=3
c
Filter=4
d
Filter=5
MCF7
MCF7M1
MCF7TR
PC1
PC2
PC3

**Extended Data Fig. 6. Filtering the cells with less contacts for clustering.**

**a**, Removing cells with the minimal contacts of 2 in 1Mb bins. **b**, Removing cells with the minimal contacts of 3 in 1Mb bins. **c**, Removing cells with the minimal contacts of 4 in 1Mb bins. **d**, Removing cells with the minimal contacts of 5 in 1Mb bins. PC1 is the first eigenvector. PC2 is the second eigenvector. PC3 is the third eigenvector.

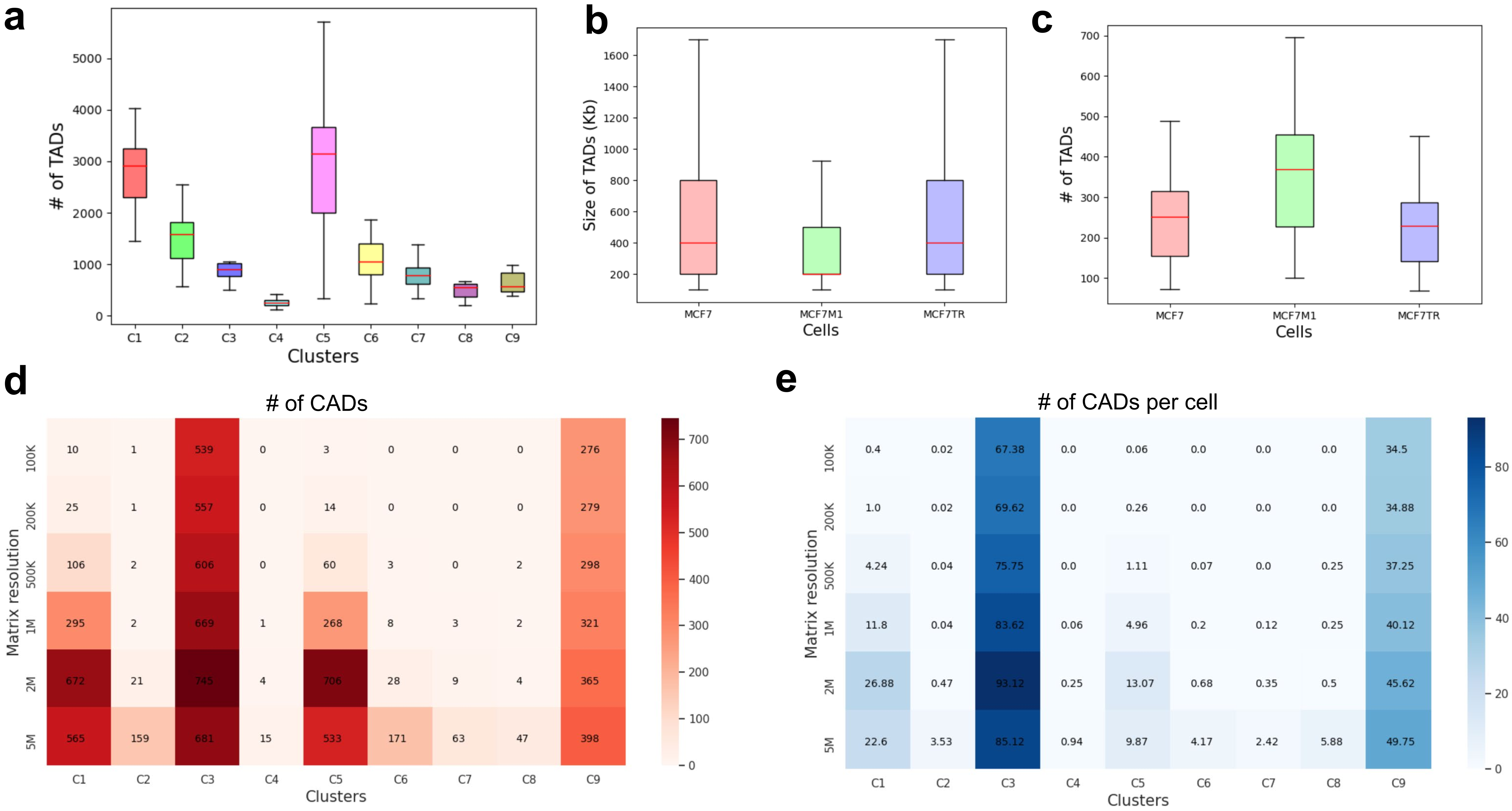

a
### of TADs
0
1000
2000
3000
4000
5000
C1 C2 C3 C4 C5 C6 C7 C8 C9
Clusters
b
Size of TADs (Kb)
200
400
600
800
1000
1200
1400
1600
MCF7 MCF7M1 MCF7TR
Cells
c
### of TADs
100
200
300
400
500
600
700
MCF7 MCF7M1 MCF7TR
Cells
d
### of CADs
Matrix resolution
100K 10 1 539 0 3 0 0 0 276
200K 25 1 557 0 14 0 0 0 279
500K 106 2 606 0 60 3 0 2 298
1M 295 2 669 1 268 8 3 2 321
2M 672 21 745 4 706 28 9 4 365
5M 565 159 681 15 533 171 63 47 398
C1 C2 C3 C4 C5 C6 C7 C8 C9
Clusters
0
100
200
300
400
500
600
700
e
### of CADs per cell
Matrix resolution
100K 0.4 0.02 67.38 0.0 0.06 0.0 0.0 0.0 34.5
200K 1.0 0.02 69.62 0.0 0.26 0.0 0.0 0.0 34.88
500K 4.24 0.04 75.75 0.0 1.11 0.07 0.0 0.25 37.25
1M 11.8 0.04 83.62 0.06 4.96 0.2 0.12 0.25 40.12
2M 26.88 0.47 93.12 0.25 13.07 0.68 0.35 0.5 45.62
5M 22.6 3.53 85.12 0.94 9.87 4.17 2.42 5.88 49.75
C1 C2 C3 C4 C5 C6 C7 C8 C9
Clusters
0
20
40
60
80

**Extended Data Fig. 7. TADs and CADs of scHi-C clusters.**

**a**, Number of TADs in each of nine clusters. **b**, Size of TADs of cancer single cells. **c**, Number of TADs of cancer single cells. **d**, Number of CADs of clusters in the matrix resolution at 100K, 200K, 500K, 1M, 2M and 5M. **e**, Number of CADs per cell of clusters in the matrix resolution at 100K, 200K, 500K, 1M, 2M and 5M.

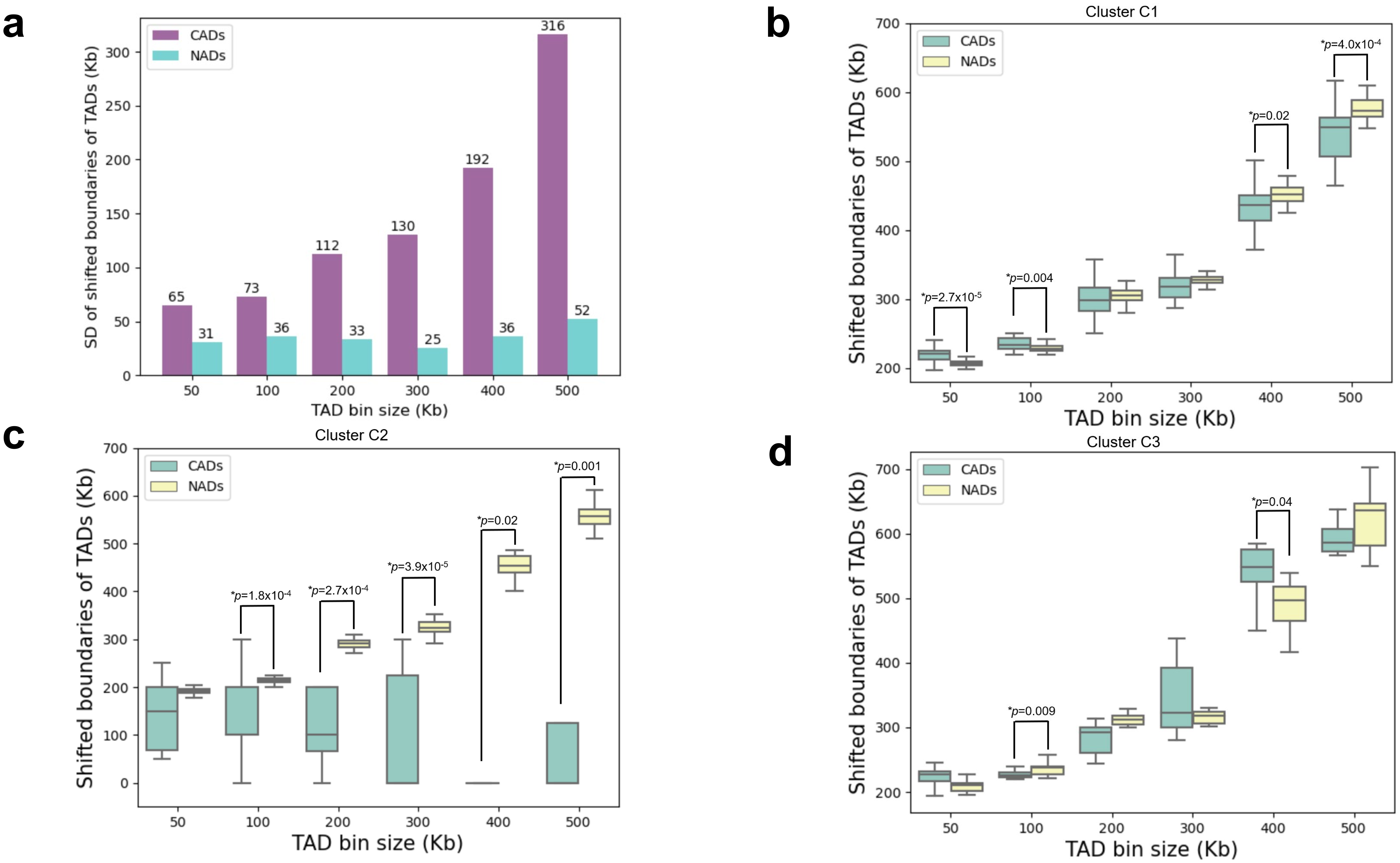

a
CADs
NADs
SD of shifted boundaries of TADs (Kb)
TAD bin size (Kb)
65 31 73 36 112 33 130 25 192 36 316 52
b
Cluster C1
Shifted boundaries of TADs (Kb)
TAD bin size (Kb)
*p=2.7x10-5
*p=0.004
*p=0.02
*p=4.0x10-4
c
Cluster C2
*p=1.8x10-4
*p=2.7x10-4
*p=3.9x10-5
*p=0.02
*p=0.001
d
Cluster C3
*p=0.009
*p=0.04

**Extended Data Fig. 8. The correlations of TADs with CADs and NADs.**

**a**, Standard deviation (SD) of shifted boundaries of TADs in CADs/NADs at the TAD bin size of 50Kb, 100Kb, 200Kb, 300Kb, 400Kb and 500Kb. **b-d**, The shifted boundaries of TADs of CADs/NADs of C1-3 at the TAD bin size of 50Kb, 100Kb, 200Kb, 300Kb, 400Kb and 500Kb, respectively. (**b**) C1, (**c**) C2, and (**d**) C3. *: Wilcoxon rank-sum test.

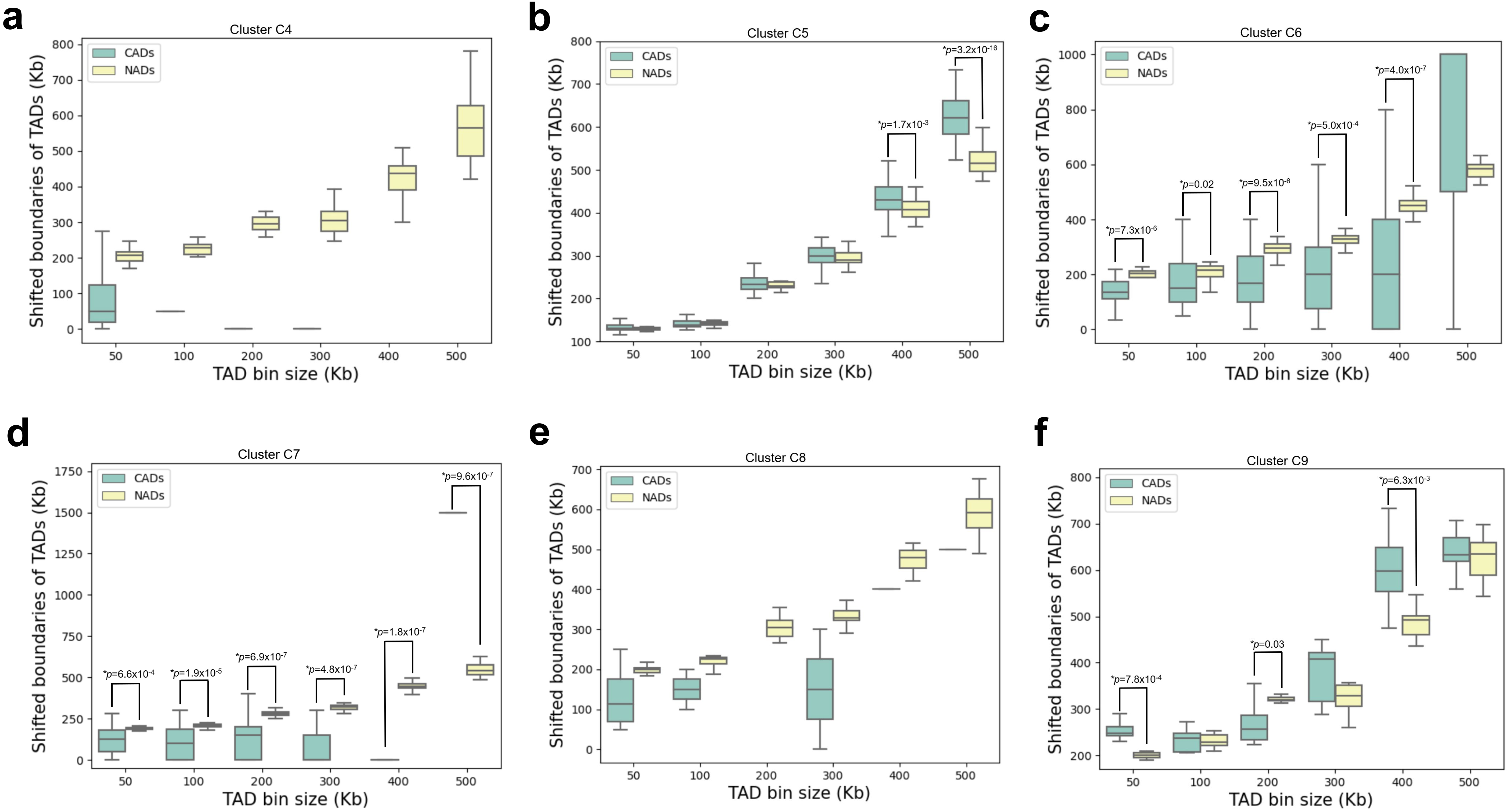

a
Cluster C4
CADs
NADs
Shifted boundaries of TADs (Kb)
TAD bin size (Kb)
b
Cluster C5
CADs
NADs
*p=1.7x10-3
*p=3.2x10-16
Shifted boundaries of TADs (Kb)
TAD bin size (Kb)
c
Cluster C6
CADs
NADs
*p=7.3x10-6
*p=0.02
*p=9.5x10-6
*p=5.0x10-4
*p=4.0x10-7
Shifted boundaries of TADs (Kb)
TAD bin size (Kb)
d
Cluster C7
CADs
NADs
*p=6.6x10-4
*p=1.9x10-5
*p=6.9x10-7
*p=4.8x10-7
*p=1.8x10-7
*p=9.6x10-7
Shifted boundaries of TADs (Kb)
TAD bin size (Kb)
e
Cluster C8
CADs
NADs
Shifted boundaries of TADs (Kb)
TAD bin size (Kb)
f
Cluster C9
CADs
NADs
*p=7.8x10-4
*p=0.03
*p=6.3x10-3
Shifted boundaries of TADs (Kb)
TAD bin size (Kb)

**Extended Data Fig. 9. The correlations of TADs with CADs and NADs in individual clusters.**

**a-f**, The shifted boundaries of TADs of CADs/NADs of C4-9 at the TAD bin size of 50Kb, 100Kb, 200Kb, 300Kb, 400Kb and 500Kb, respectively. (**a**) C4, (**b**) C5, (**c**) C6, (**d**) C7, (**e**) C8 and (**f**) C9. *: Wilcoxon rank-sum test.

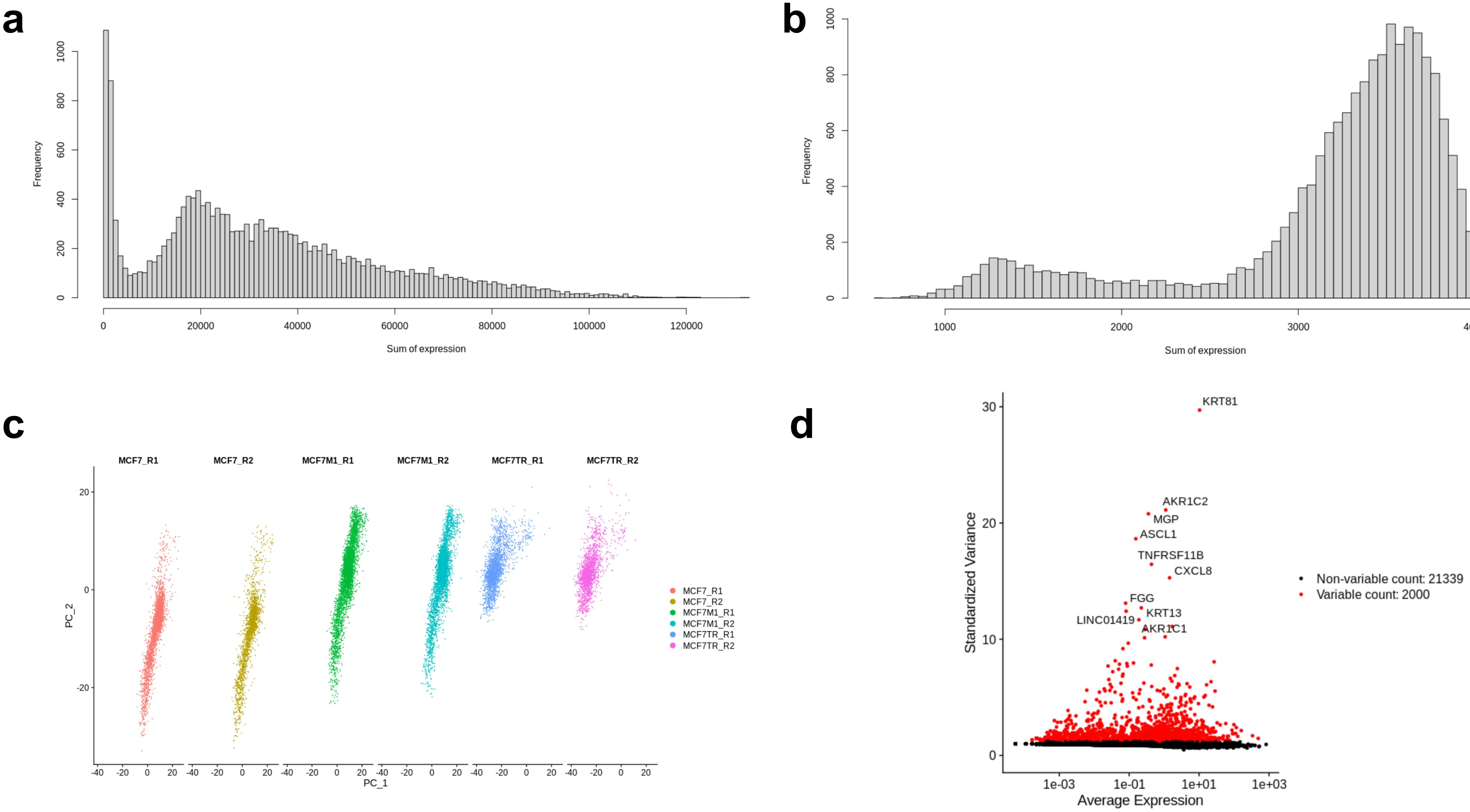

a
Frequency
Sum of expression
b
Frequency
Sum of expression
c
MCF7_R1
MCF7_R2
MCF7M1_R1
MCF7M1_R2
MCF7TR_R1
MCF7TR_R2
PC_2
PC_1
d
KRT81
AKR1C2
MGP
ASCL1
TNFRSF11B
CXCL8
FGG
KRT13
LINC01419
AKR1C1
Standardized Variance
Average Expression
Non-variable count: 21339
Variable count: 2000

**Extended Data Fig. 10. Summary of scRNA-seq data.**

**a**, The total gene expression frequency of individual cells before normalization. **b**, The total gene expression frequency of individual cells after normalization. **c**, The replicates of scRNA-seq data. PC_1 is the first eigenvector and PC_2 is the second eigenvector. **d**, The standardized variance of genes. Top 2000 genes were colored with red and top 10 genes were additionally labelled with their gene symbols.

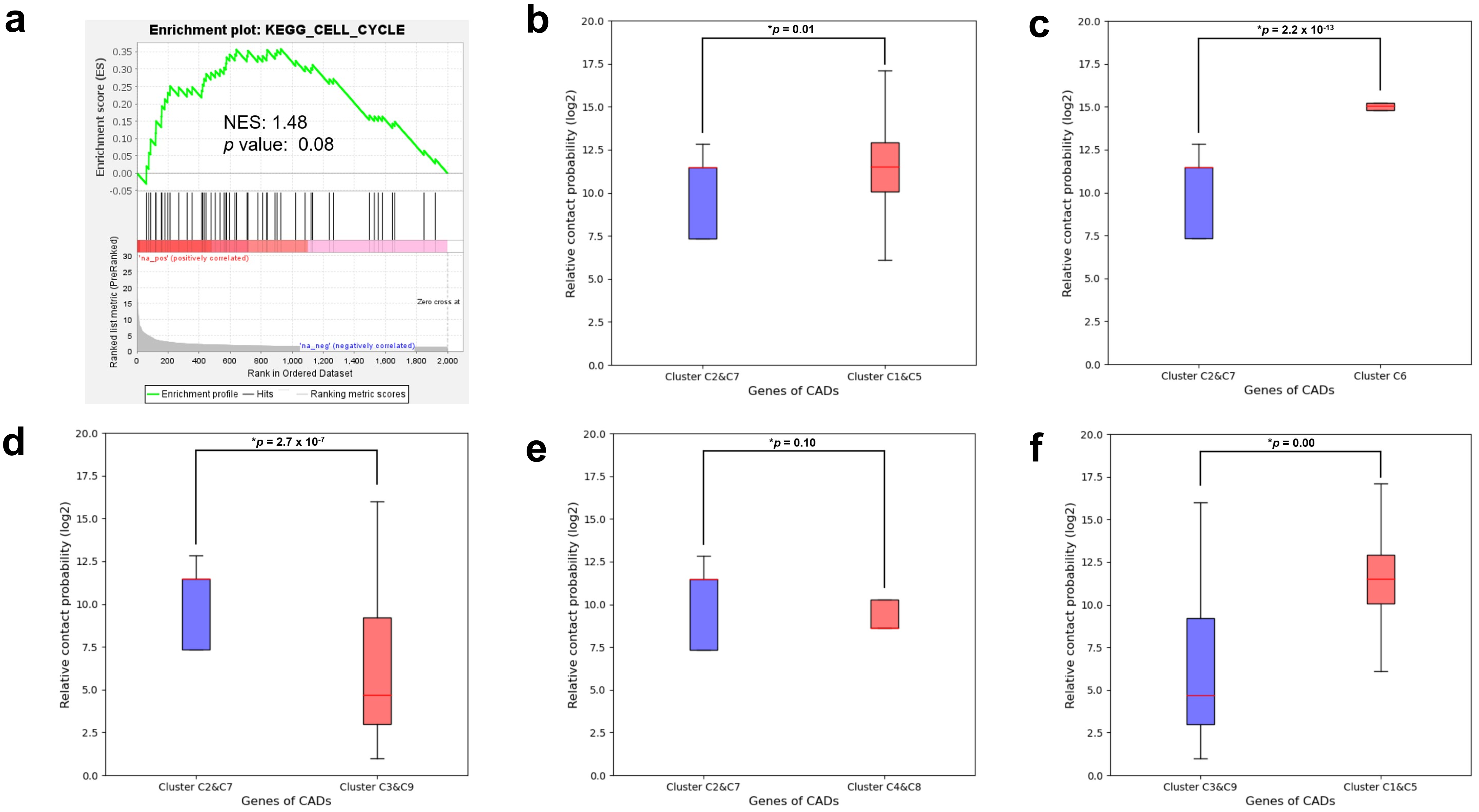

a
Enrichment plot: KEGG_CELL_CYCLE
Enrichment score (ES)
NES: 1.48
p value: 0.08
'na_pos' (positively correlated)
Zero cross at
'na_neg' (negatively correlated)
Ranked list metric (PreRanked)
Rank in Ordered Dataset
Enrichment profile
Hits
Ranking metric scores
b
*p = 0.01
Relative contact probability (log2)
Cluster C2&C7
Cluster C1&C5
Genes of CADs
c
*p = 2.2 x 10-13
Cluster C2&C7
Cluster C6
Genes of CADs
d
*p = 2.7 x 10-7
Cluster C2&C7
Cluster C3&C9
Genes of CADs
e
*p = 0.10
Cluster C2&C7
Cluster C4&C8
Genes of CADs
f
*p = 0.00
Cluster C3&C9
Cluster C1&C5
Genes of CADs

**Extended Data Fig. 11. The enrichment and relative contact probability of genes.**

**a**, Enrichment of KEGG cell cycle signaling pathway for top 2000 variable genes in scRNA-seq clusters. NES: Normalized Enrichment Score. **b-f**, The comparison of relative contact probability of genes of CADs in the combination of scHi-C clusters. (**b**) C2&C7 vs C1&C5. (**c**) C2&C7 vs C6. (**d**) C2&C7 vs C3&C9. (**e**) C2&C7 vs C4&C8. (**f**) C3&C9 vs C1&C5. *: Wilcoxon rank-sum test.

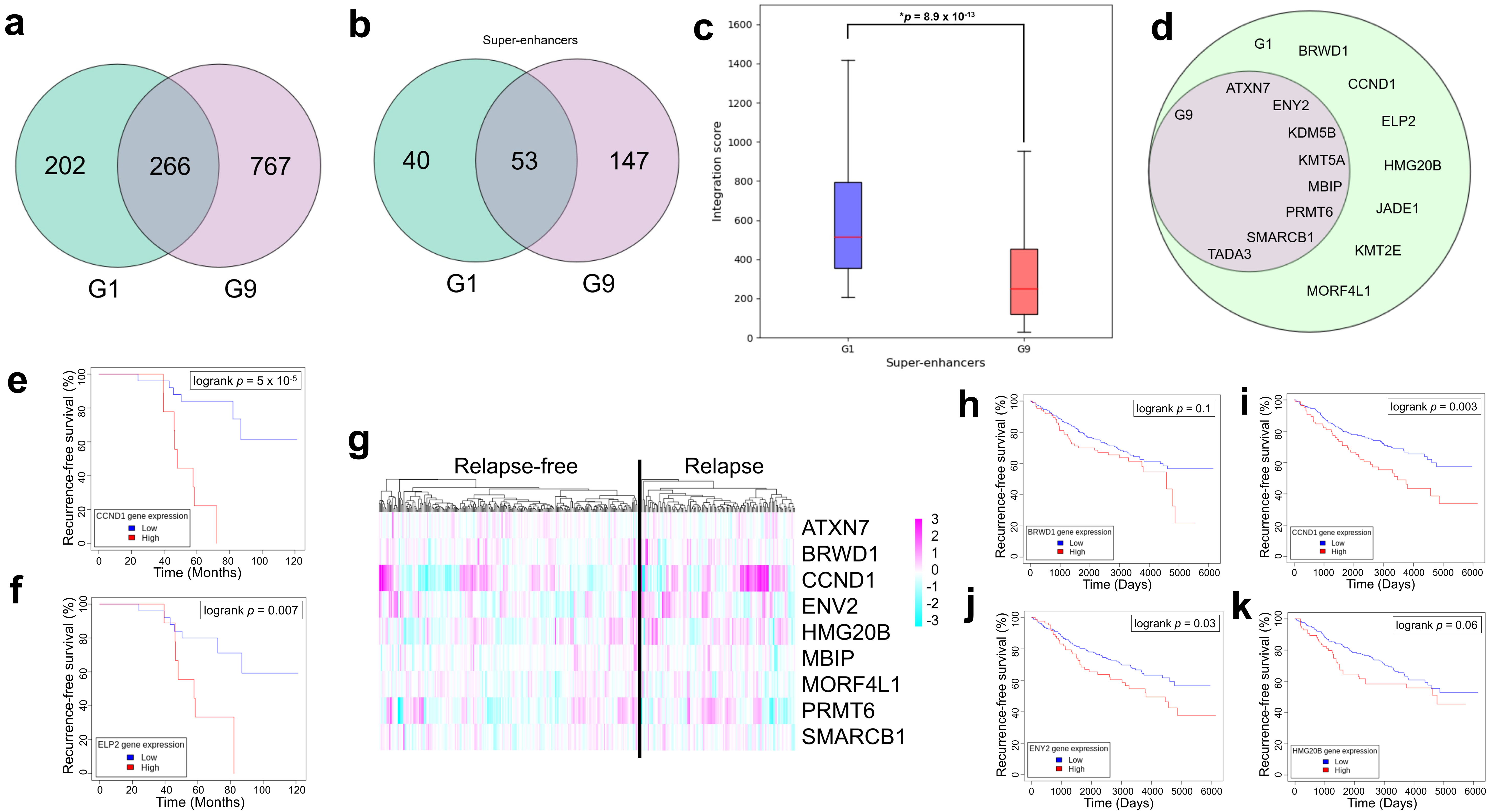
a
202
266
767
G1
G9
b
Super-enhancers
40
53
147
G1
G9
c
*p = 8.9 x 10^-13
Integration score
Super-enhancers
G1
G9
d
G1
G9
BRWD1
CCND1
ELP2
HMG20B
JADE1
KMT2E
MORF4L1
ATXN7
ENY2
KDM5B
KMT5A
MBIP
PRMT6
SMARCB1
TADA3
e
logrank p = 5 x 10-5
Recurrence-free survival (%)
Time (Months)
CCND1 gene expression
Low
High
f
logrank p = 0.007
Recurrence-free survival (%)
Time (Months)
ELP2 gene expression
Low
High
g
Relapse-free
Relapse
ATXN7
BRWD1
CCND1
ENV2
HMG20B
MBIP
MORF4L1
PRMT6
SMARCB1
h
logrank p = 0.1
Recurrence-free survival (%)
Time (Days)
BRWD1 gene expression
Low
High
i
logrank p = 0.003
Recurrence-free survival (%)
Time (Days)
CCND1 gene expression
Low
High
j
logrank p = 0.03
Recurrence-free survival (%)
Time (Days)
ENY2 gene expression
Low
High
k
logrank p = 0.06
Recurrence-free survival (%)
Time (Days)
HMG20B gene expression
Low
High

**Extended Data Fig. 12. The integration score and survival analysis of chromatin modifying enzymes in TISPs.**

**a**, The number of genes in G1 and G9. **b**, The genes marked with super-enhancers in G1 and G9. **c**, The integration score of G1 and G9 super-enhancers. **d**, List of 15 chromatin modifying enzymes involved in G1, and G9. **e-f**, Recurrence-free survival of chromatin modifying enzymes in cohort GSE2990. (e) CCND1. (f) ELP2. The patients were stratified by gene expression levels at the top quartile (25%) vs the rest (75%). p value was determined by the log-rank test. **g**, The expression of chromatin modifying enzymes in relapse-free and relapse breast cancer patient cohort GSE6532. **h-k**, Recurrence-free survival of chromatin modifying enzymes in cohort GSE6532. (**h**) BRWD1. (**i**) CCND1. (**j**) ENY2. (**k**) HMG20B. The patients were stratified by gene expression levels at the top quartile (25%) vs the rest (75%). *p* value was determined by the log-rank test.

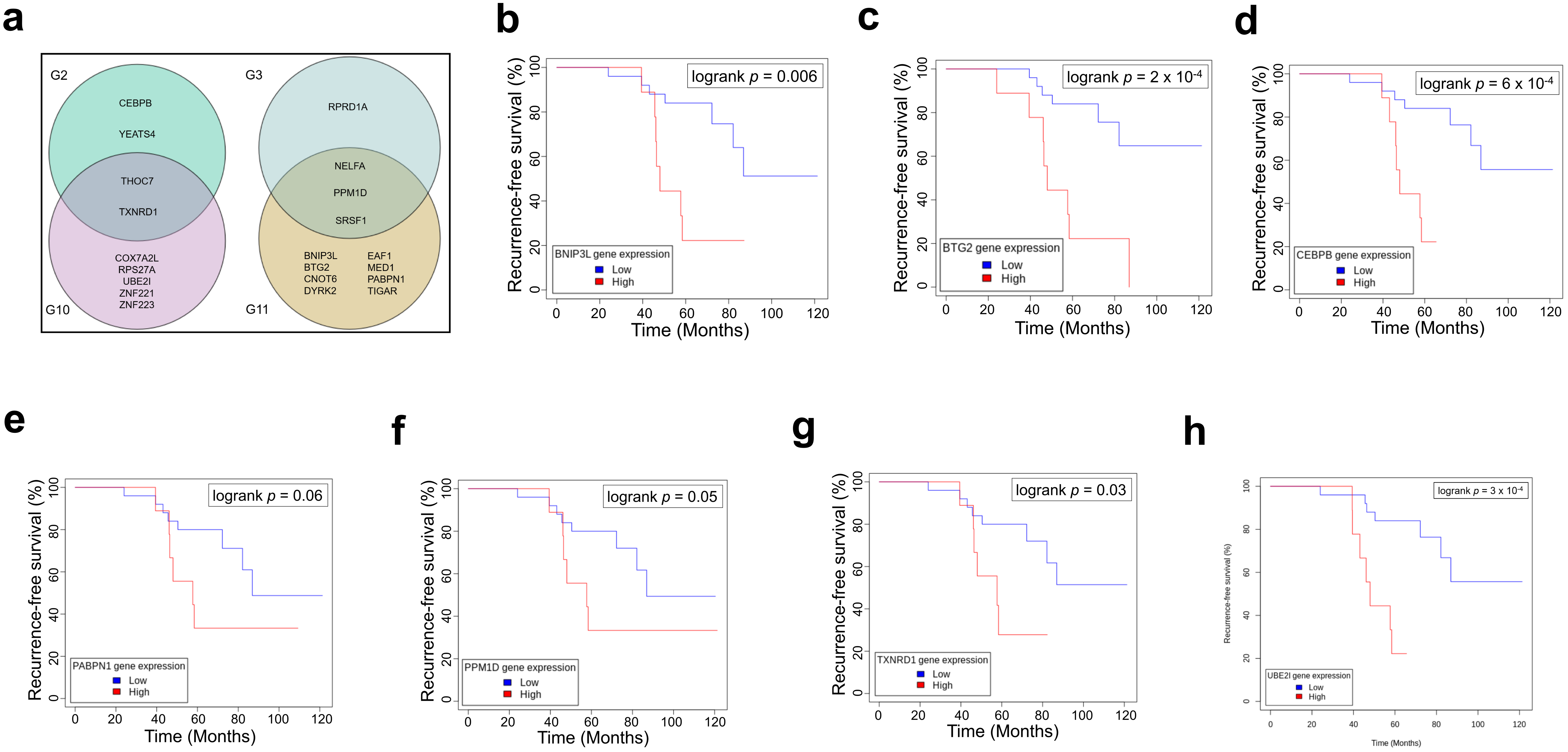
a
G2
G3
G10
G11
CEBPB
YEATS4
THOC7
TXNRD1
COX7A2L
RPS27A
UBE2I
ZNF221
ZNF223
RPRD1A
NELFA
PPM1D
SRSF1
BNIP3L
BTG2
CNOT6
DYRK2
EAF1
MED1
PABPN1
TIGAR
b
logrank p = 0.006
BNIP3L gene expression
Low
High
c
logrank p = 2 x 10-4
BTG2 gene expression
Low
High
d
logrank p = 6 x 10-4
CEBPB gene expression
Low
High
e
logrank p = 0.06
PABPN1 gene expression
Low
High
f
logrank p = 0.05
PPM1D gene expression
Low
High
g
logrank p = 0.03
TXNRD1 gene expression
Low
High
h
logrank p = 3 x 10-4
UBE2I gene expression
Low
High
Recurrence-free survival (%)
Time (Months)

**Extended Data Fig. 13. The survival analysis of transcription regulators in breast cancer patient cohorts.**

**a**, List of 21 transcription regulators involved in G2, G3, G10 and G11. **b-f,** Recurrence-free survival of transcription regulators in cohort GSE2990. (**b**) BNIP3L. (**c**) BTG2. (**d**) CEBPB. (**e**) PABPN1. (**f**) PPM1D. (**g**) TXNRD1. (**h**) UBE2I. The patients were stratified by gene expression levels at the top quartile (25%) vs the rest (75%). *p* value was determined by the log-rank test.

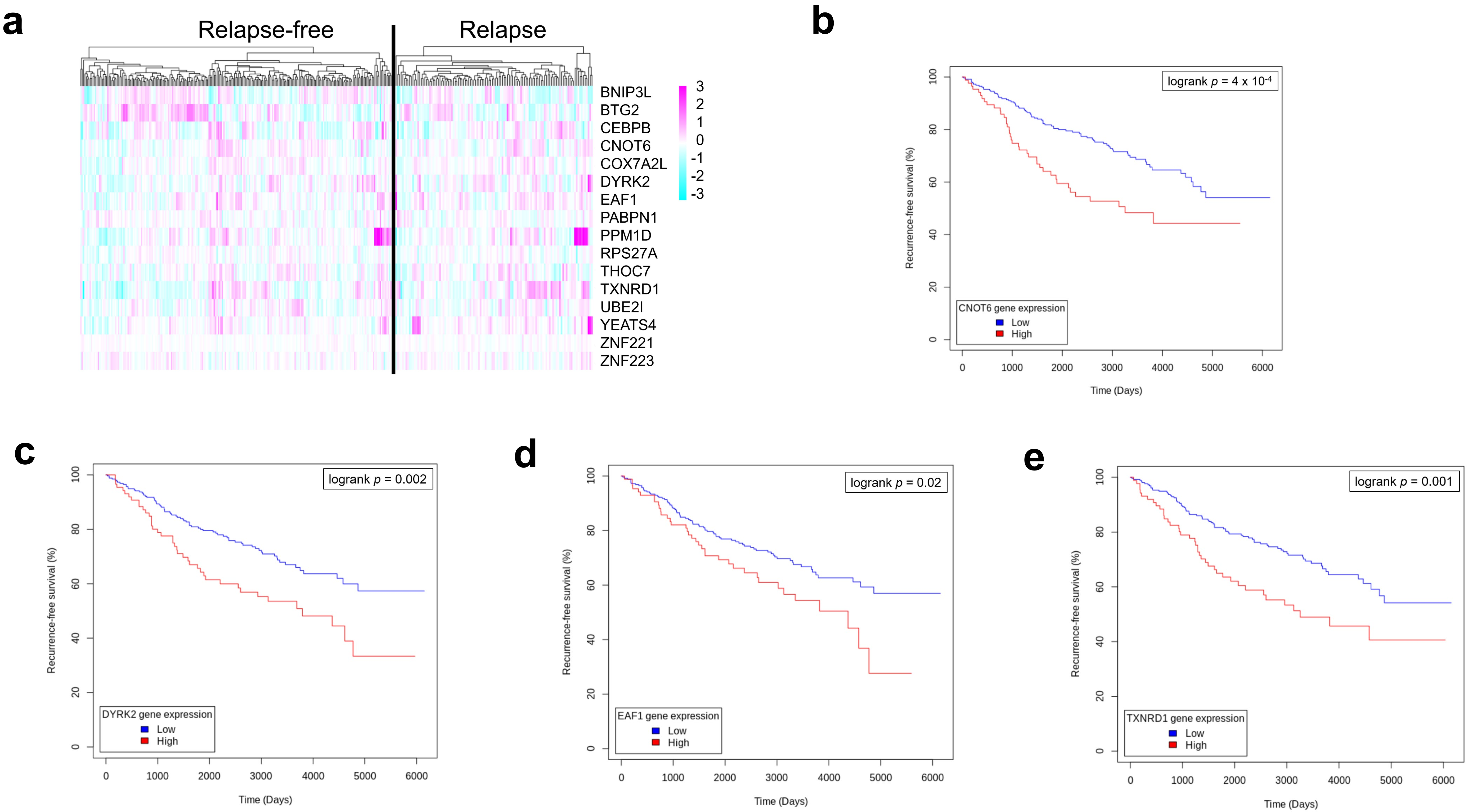

a
Relapse-free
Relapse
BNIP3L
BTG2
CEBPB
CNOT6
COX7A2L
DYRK2
EAF1
PABPN1
PPM1D
RPS27A
THOC7
TXNRD1
UBE2I
YEATS4
ZNF221
ZNF223
3
2
1
0
-1
-2
-3
b
logrank p = 4 x 10-4
CNOT6 gene expression
Low
High
Recurrence-free survival (%)
Time (Days)
c
logrank p = 0.002
DYRK2 gene expression
Low
High
Recurrence-free survival (%)
Time (Days)
d
logrank p = 0.02
EAF1 gene expression
Low
High
Recurrence-free survival (%)
Time (Days)
e
logrank p = 0.001
TXNRD1 gene expression
Low
High
Recurrence-free survival (%)
Time (Days)

**Extended Data Fig. 14. The expression and survival analysis of transcription regulators in breast cancer patient cohorts.**

**a**, The expression of transcription regulators in relapse-free and relapse breast cancer patient cohort GSE6532. **b-e**, Recurrence-free survival of transcription regulators in cohort GSE6532. (**b**) CNOT6. (**c**) DYRK2. (**d**) EAF1 (**e**) TXNRD1. The patients were stratified by gene expression levels at the top quartile (25%) vs the rest (75%). *p* value was determined by the log-rank test.

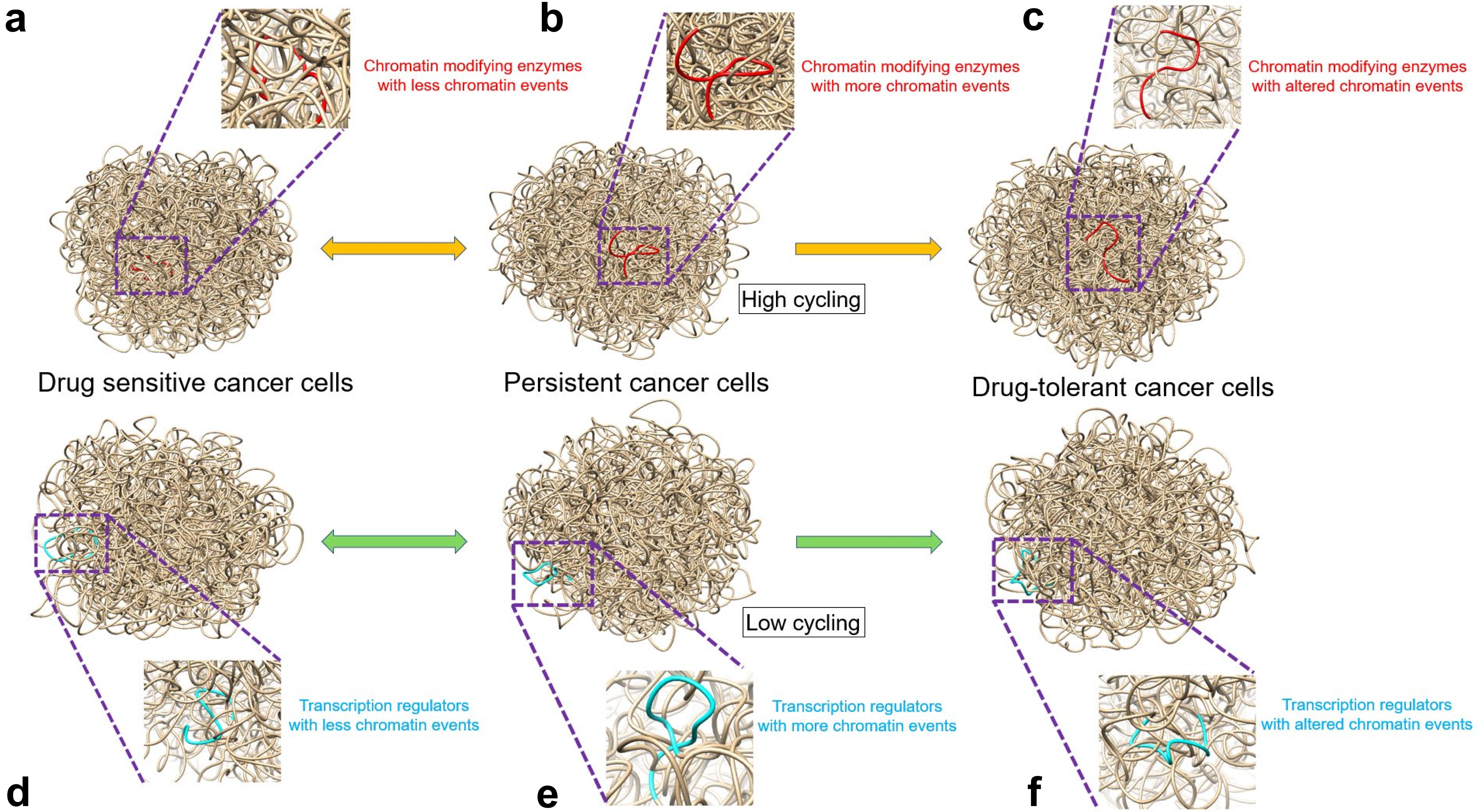
a
Chromatin modifying enzymes with less chromatin events
b
Chromatin modifying enzymes with more chromatin events
c
Chromatin modifying enzymes with altered chromatin events
High cycling
Drug sensitive cancer cells
Persistent cancer cells
Drug-tolerant cancer cells
Low cycling
Transcription regulators with less chromatin events
Transcription regulators with more chromatin events
Transcription regulators with altered chromatin events
d
e
f

**Extended Data Fig. 15. A proposed model for regulating 3D chromatin structures of drug-tolerant cancer cells by two distinctive pathways.**

**a**, Drug-sensitive cancer cells with lower expression of chromatin modifying enzymes. **b**, Cycling persistent cancer cells with higher expression of chromatin modifying enzymes. **c**, Cycling drug-tolerant cancer cells. **d**, Drug-sensitive cancer cells with lower expression of transcription regulators. **e**, Non-cycling persistent cancer cells with higher expression of transcription regulators. **f**, Non-cycling drug-tolerant cancer cells. **a-c**, an example of chromatin modifying enzyme, JADE1 labelled with red in the brown background of chromosome 4. **d-f**, an example of transcription regulator, TXNRD1 labelled with cyan in the brown background of chromosome 12.
